# Revisiting fungal polarized growth: from spitzenkörper to crescent accumulation of secretory vesicles

**DOI:** 10.1101/2024.12.25.629896

**Authors:** Adrien Hamandjian, Glen Calvar, Matthieu Blandenet, Mélanie Crumière, Nathalie Poussereau, Mathias Choquer, Christophe Bruel

## Abstract

Polarized elongation in filamentous fungi depends on secretory vesicles being supplied and fusing at the apex. In Ascomycota and Basidiomycota, these vesicles have mostly been reported to accumulate into a spheroidal structure known as the spitzenkörper. Using time-lapse microscopy, the spatial and temporal dynamics of fluorescently-labelled vesicles were investigated in 13 apices of the phytopathogenic fungus *Botrytis cinerea*. Over time, the fluorescent signal highlighted the spheroid spitzenkörper in half the sample and a crescent-shaped region of the apical dome in the other half. A linear relationship was found between the roundness of the fluorescent region and the hyphal elongation rate. Temporal Dynamics Clustering and Fourier transform showed periodic pulses of fluorescence intensity in the spitzenkörper that were absent in the crescent-shape region. These results reveal a dual mode of secretory vesicles accumulation at the apex of growing hyphae.

**Highlights:**

- Secretory Vesicles accumulation at the apex form a spheroid spitzenkörper or a crescent
- Vesicles accumulation of at the spitzenkörper is periodic
- Roundness of SV accumulation regions correlates with elongation rates

## Introduction

Sustained polar cell elongation is a shared trait in various branches of the tree of life, from filamentous bacteria such as actinomycetes^1^, to tip-growing eukaryotic cells found in plants^2–4^ (e.g., pollen tubes and root hairs) and algae^5^ (e.g., protonemata). In eukaryotic microorganisms, it has been described in filamentous yeasts, including pathogens such as *Candida albicans*^6,7^ or *Ashbya gossypii*^8^ whose respective virulence to human and cotton relies on the production of filaments. More notably, polar cell elongation is a hallmark of filamentous fungi, supporting continuous hyphal extension.

Polarized cell elongation results from the spatial restriction of exocytosis within the fungal cell. By limiting vesicle fusion to a specific region, lipids are locally supplied to the plasma membrane, resulting in the unidirectional elongation of the cell^9^. The development of the mycelium, formed by extending and branching hyphae, therefore depends on the production, transport, and regulated exocytosis of secretory vesicles (SV) at the hyphal tip. SV are Golgi-derived vesicles transported to the apex by microtubules^10–12^ or directly produced on-site by apical Golgi compartments^13^. Once at the hyphal tip, also called the apex, SV are locally transported along actin filaments to interact with the exocyst, a protein complex that facilitates vesicle docking and fusion with the plasma membrane during exocytosis^14–16^.

In addition to transporting and releasing lipids, SV also carry transmembrane proteins destined for the plasma membrane, and soluble proteins delivered to the extracellular space, including the cell wall. Among the transmembrane protein cargoes, chitin synthases (CHS) have received particular attention. CHS are type 2 glycosyltransferases that synthesize chitin, a major structural component of the fungal cell wall that is composed of N-acetylglucosamine residues linked by beta-(1,4) linkages^17^. Because CHS are transported by SV to the apical plasma membrane, fluorescently tagged CHS have been widely used to visualize SV trafficking^11,18–20^. The delivery by SV of enzymes involved in cell wall synthesis and remodeling ensures that cell wall expansion occurs in coordination with plasma membrane extension during apical elongation.

SV are the final carriers of all proteins that follow the canonical secretory pathway, i.e. proteins translated in the endoplasmic reticulum and matured in the Golgi apparatus^21^. These proteins include lytic enzymes involved in fungal nutrition, which play a key role in ecosystems due to their importance in nutrient cycling^22,23^. Additionally, SV transport proteins that mediate interactions between fungi and other organisms, notably virulence factors secreted by pathogenic species^24^. Beyond its ecological function, fungal secretion is also crucial to various industrial processes, ranging from fermentation in the food industry to the secretion of proteins and enzymes of economic interest^25^. Consequently, a deeper understanding of temporal and spatial dynamics of exocytosis in fungi is needed to better understand their biology, but also to better control pathogens or, on the contrary, maximize production yields in industrial settings.

In Ascomycota and Basidiomycota, the apex of elongating filaments is marked by a visible accumulation of vesicles forming a spheroidal structure known as the spitzenkörper^26^. This structure has been proposed to act as a vesicle supply center (VSC), where vesicles accumulate before being delivered to the plasma membrane^27^. The spitzenkörper is located at the interface between the microtubules transporting SV to the apex^28^, the polarisome nucleating actin filaments at the apex^29^, and the exocyst mediating the vesicle tethering and fusion with the plasma membrane^30^. In *Neurospora crassa*, it was shown to be stratified with SV subpopulations known as microvesicles and macrovesicles accumulating at the core and in the outer layer of the spitzenkörper, respectively^31^.

The spitzenkörper is therefore an organized accumulation of SV at the apex which can be described as both dynamic and stable. It is dynamic by the constant entry and exit of vesicles^26,32–34^, but stable by its importance to direct growth in actively growing filaments^35–38^. This stability is however challenged. In the filament-producing yeast *Ashbya gossypii*, chemical labelling of lipids showed the presence of a spitzenkörper in fast elongating hyphae, but a crescent-shaped accumulation of vesicles in slower elongating hyphae^39^. Interestingly, both spitzenkörper and crescent-shaped accumulation of SV have also been documented in *Aspergillus nidulans*^19,40^. While significant progress has been made in understanding the apical machinery and vesicle trafficking during polarized cell elongation, the consistency of vesicle accumulation through time and space across filaments remains unclear. In addition, studies on polarized cell elongation have mainly focused on cells displaying a spitzenkörper and have been conducted in saprotrophic model species, potentially underestimating the diversity of vesicle accumulation patterns and behaviors both between hyphae and across fungal species.

In this study, a fluorescently-labelled class III chitin synthase was used to label SV in the phytopathogenic necrotrophic fungus *Botrytis cinerea*. Time-lapse confocal acquisitions revealed two populations of growing hyphae displaying either a spheroid spitzenkörper or a fluorescent crescent at the apex. Further temporal analysis revealed periodic variations of fluorescence intensity at the apex of hyphae displaying a spitzenkörper, that were not detected in hyphae displaying a crescent. These results reveal the existence of at least two populations of hyphae that display different spatial and temporal dynamics of SV distribution in the apical dome.

## Results

### 1. The fluorescently-labelled class III chitin synthase BcCHSIIIa localizes at the tip of growing hyphae in *B. cinerea*

To visualize SV at the apex of *B. cinerea* hyphae, a reporter system was required. Based on the successful labelling of SV through the fluorescent tagging of class III chitin synthases in different ascomycetes^11,18–20,40^, the *B. cinerea* chitin synthase IIIa (BcCHSIIIa - Bcin04g03120) was chosen^41^. The gene encoding an optimized version of the fluorescent protein eGFP^42^ was fused at the 3’ end of the BcCHSIIIa gene by homologous recombination to produce the chimeric BcCHSIIIa::eGFP protein (Fig.S1). In parallel, a control strain was constructed that produces that eGFP alone under the control of the constitutive pOliC promoter (eGFP^Cyt^ strain).

Following growth on a glass surface, confocal microscopy was performed on both strains. As expected, the eGFP^Cyt^ control strain displayed a diffuse cytosolic fluorescent signal (Fig.1A). In contrast, mobile fluorescent speckles and an apical accumulation of fluorescence were observed in the BcCHSIIIa::eGFP strain (Fig.1B) that were consistent with previously reported localizations of SV transporting chitin synthases^18–20,40^. These observations therefore validated the creation of the desired reporter strain.

**Figure 1.**
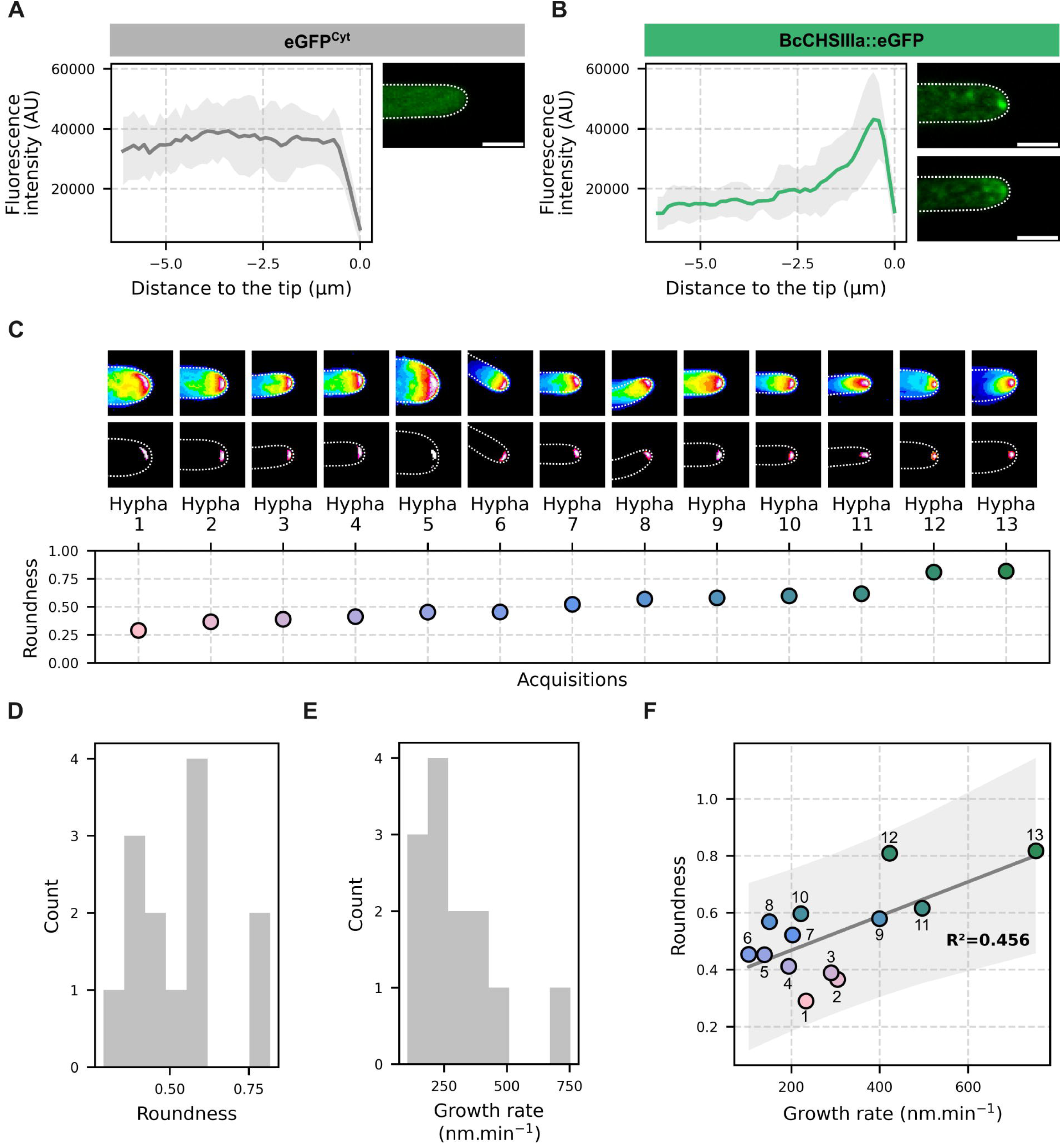
Distinct accumulation patterns of BcCHSIIIa::eGFP correlate with hyphal growth rates. (A–B) Fluorescence intensity measured along the longitudinal axis of growing hyphae from the eGFP^Cyt^ strain (A) and the BcCHSIIIa::eGFP strain (B). Solid lines represent mean fluorescence intensity, shaded areas indicate standard deviation (n = 5 for A, n = 12 for B). Confocal images show representative hyphae (scale bar = 3 µm). (C, top) Time projections after apex tracking, showing the overall fluorescence distribution over the acquisition time. (C, middle) Region of fluorescence accumulation, defined by the pixels with the top 1% intensity in (C, top). Dotted white lines outline hyphae. Regions of interest displayed in (C, top) and (C, middle) are 6.78x6.78 µm. (C, bottom) Scatter plot of roundness values measured from regions of accumulation in (C, middle). Colors indicate roundness values. (D) The distribution of roundness measures follows a normal distribution (Shapiro-Wilk test, *p*-value 0.446). (E) The distribution of growth rate measures also follows a normal distribution (Shapiro-Wilk test, *p*-value 0.063). (F) Scatter plot of roundness versus growth rate for each acquisition. The solid grey line shows the linear regression, the grey-shaded area represents the 95% prediction interval for individual observations. Data points are color-coded according to roundness values as in (C). Numbers in (F) correspond to Hypha 1 to 13 as defined in (C).

### 2. Patterns of apical BcCHSIIIa::eGFP accumulation are linked to hyphal elongation rate

Noticeably, in different hyphae of the BcCHSIIIa::eGFP strain, the fluorescence signal at the apex appeared either as a spheroid spot or was more diffused (Fig.1B, right panel top and bottom). To exclude the possibility of an artifact, 10 min time-lapse acquisitions (1 frame every 4.455 sec) were collected on 13 BcCHSIIIa::eGFP hyphae (Fig.1C). For each acquisition, the apex was tracked and cropped to accommodate for hyphal elongation. The overall distribution of fluorescence over the course of the acquisition was visualized by performing time-projections (Fig. 1C, top). The region of maximal fluorescence accumulation (see *Material and methods*) was then extracted for each hypha using a threshold-based approach (Fig.1C middle). In some hyphae, this region showed as a spheroid spot, reflecting a spatially confined accumulation of the fluorescent SV over time. In other hyphae it showed as a crescent. This crescent was not necessarily occupied at once, but resulted from changes over time in the position of the fluorescent signal that shaped a crescent when cumulating all the frames in a projection. The area of both the spheroid and crescent-shaped regions of fluorescence accumulation were measured and followed a normal distribution (Shapiro-Wilk test, *p*-value 0.297, mean area of 0.428 ± 0.019 µm²). The roundness of these regions were also measured and were normally distributed, ranging between 0.290 and 0.818 (Shapiro-Wilk test, *p*-value 0.446, mean roundness of 0.529 ± 0.154). This last observations indicate that the apical region of fluorescence accumulation spans a continuum of shapes, ranging from crescent-like to spherical.

For each acquisition, kymographs were then produced to estimate the growth rate of each hyphae during acquisition (Fig.S2). This rate varied between 104 and 753 nm.min^-1^ (mean value 301 ± 173 nm.min^-1^) and no statistical difference was observed between the control (eGFP^Cyt^) and the BcCHIIIa::eGFP strain (Student t-test, *p*-value 0.190, mean growth rates of 450 ± 247 nm.min^-1^ and 301 ± 173 nm.min^-1^ with n = 5 and n = 13, respectively). In contrast, a positive linear correlation was observed between the roundness of the region of fluorescence accumulation and the hyphal elongation rate in acquisitions of the BcCHSIIIa::eGFP strain (Fig.1F, Pearson correlation 0.675, *p*-value 0.011, analysis of residues in Figure S3).

### 3. Temporal Dynamics Clustering reveals two clusters based on fluorescence variability over time

In addition to analyzing the spatial organization of SV at the apex, we examined the temporal dynamics of their accumulation. For each acquisition and at every frame, the mean intracellular fluorescence intensity was measured in the apical region (4 µm long from the tip, see *Materials and Methods*) of hyphae from the control eGFP^Cyt^ strain expressing cytoplasmic eGFP and from the BcCHSIIIa::eGFP strain. This yielded, for each acquisition, a time series representing the mean fluorescence intensity at the apex over time. In all time series this intensity fluctuated over time, independently of movements in the z dimension (Fig.S4). Since such fluctuations can result from intrinsic properties of eGFP (folding, maturation or stability), we compared the dynamics observed in the BcCHSIIIa::eGFP strain to those of the control strain expressing cytosolic eGFP.

To characterize the temporal dynamics of the fluorescence signals, we applied Temporal Dynamics Clustering (TDC)^43^, a features-based clustering approach that groups time series based on fluorescence variability over time. As drawn in figure 2A, the signals were detrended and the first-order differences were computed for each time series and expressed as a cumulative first-order difference (Fig.2B). Next, TDC parameters *u* and *Iq* were calculated to partition each cumulative distribution into two sections: the *main* section, representing common variations, and the *tail* section, capturing extreme fluctuations (Fig. 2B). For each distribution, the mean value of elements in the *main* section (µ_main_), the associated standard deviation (σ_main_), and the mean value of elements from the *tail* section (µ_tail_) were calculated. These three metrics were then used as features for *k*-means clustering, which identified two distinct clusters, termed Cluster 1 and Cluster 2 (Fig.2C).

**Figure 2.**
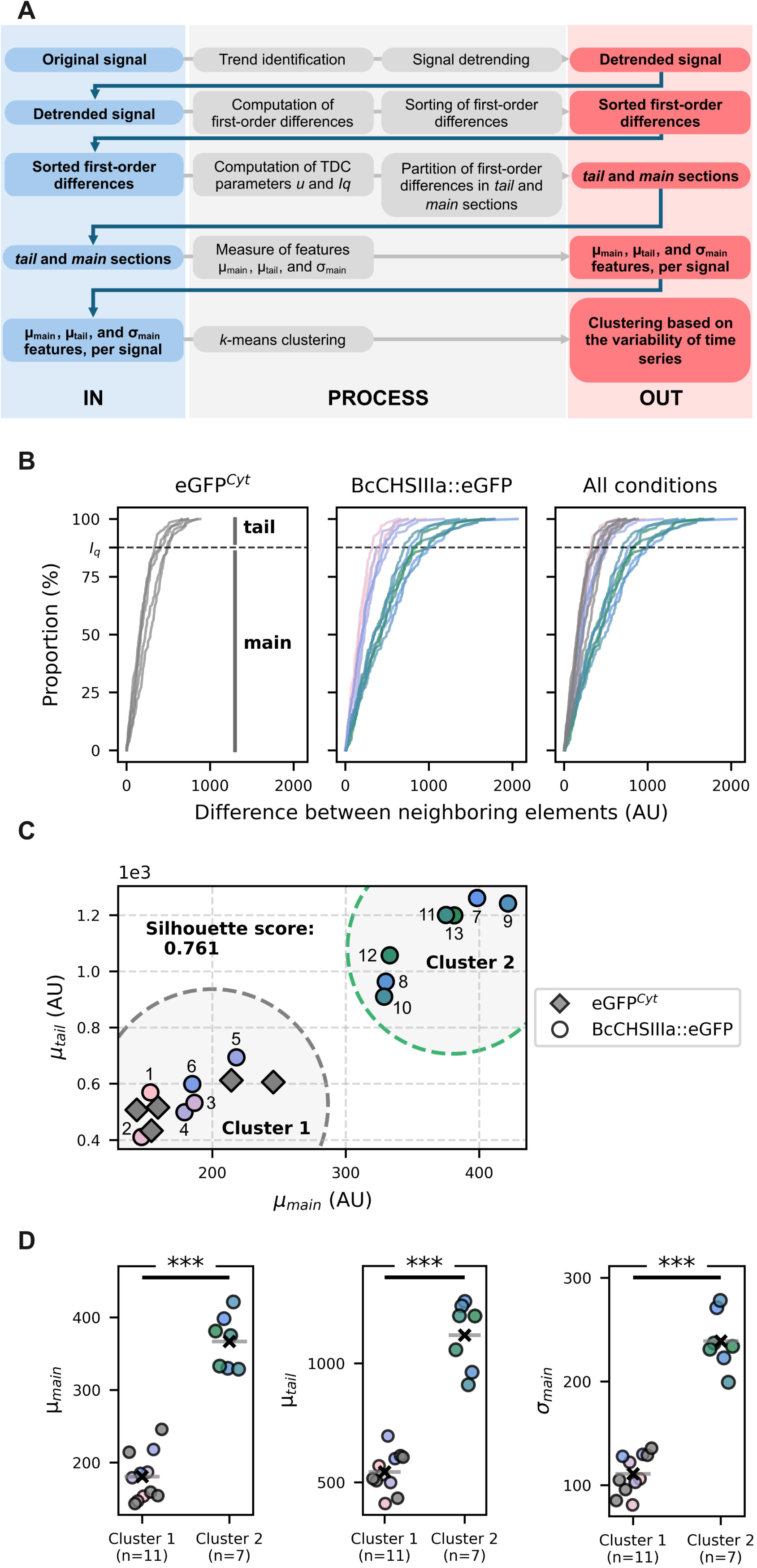
Temporal Dynamics Clustering reveals two distinct clusters differing in their variability over time. (A) Schematic overview of the Temporal Dynamics Clustering (TDC) approach. Original signals were detrended (first line), and first-order differences were computed and sorted (second line). The sorted values were partitioned into *main* and *tail* sections using the TDC-defined parameters *u* and *Iq* (third line). From each acquisition, three parameters were extracted, the mean of elements in the *main* section (µ_main_), the associated standard deviation (σ_main_) and the mean of elements in the *tail* section (µ_tail_) (fourth line). These features were used for *k*-means clustering to group signals based on fluorescence variability (fifth line). (B) Sorted first-order differences for acquisitions of the control eGFP^Cyt^ strain (left, n = 5), BcCHSIIIa::eGFP strain (middle, n = 13), or combined data (right). Dashed horizontal lines indicate the threshold *Iq* used to separate the *main* and *tail* sections. (C) Scatter plot of µ_tail_ over µ_main_ for all acquisitions of the eGFP^Cyt^ strain (diamond) and the BcCHSIIIa::eGFP strain (circles). Dashed circle outlines are used to visualize Cluster 1 and Cluster 2 identified by *k*-means. BcCHSIIIa::eGFP data points are color-coded according to the roundness of the fluorescence accumulation region as in Figure 1, and numbered to match the corresponding hyphae shown in Figure 1C. (D) Comparison of µ_main_, µ_tail_ and σ_main_ between Cluster 1 and Cluster 2. Student’s t-test *p*-values are 6.7e-9 for µ_main_, 7.7e-9 for µ_tail_ and 2.7e-9 for σ_main_.

The clustering quality was supported by a silhouette score of 0.761, indicating strong and well-defined separation between clusters. Moreover, statistical analysis revealed significant differences between clusters for all 3 TDC metrics (Fig.2D, Student’s t-test, *p*-values of 6.7e-9 for µ_main_, 2.7e-9 for σ_main_, and 7.7e-9 for µ_tail_). These differences in the 3 metrics suggest that fluorescence variability between clusters differs not only in extreme events (µ_tail_) but also in their overall temporal behavior (µ_main_). In contrast, the median fluorescence intensity before detrending was not significantly different between clusters (Mann-Whitney U test, *p-*value 0.073), suggesting that the distinction arose from differences in temporal variability rather than absolute intensity. Together, these results demonstrate the presence of two statistically distinct patterns of apical fluorescence dynamics in our sample of hyphae.

### 4. Distinct fluorescence dynamics are linked to the roundness of apical vesicle accumulation

Cluster 1 included all the acquisitions recorded on the 5 hyphae from the control eGFP^Cyt^ strain. Since clustering was based on fluorescence variability over time, Cluster 1 therefore likely reflects the intrinsic fluorescence fluctuations of eGFP itself. Interestingly, the 13 acquisitions recorded on the 13 hyphae from the BcCHSIIIa::eGFP strain distributed across both clusters (6 in Cluster 1 and 7 in Cluster 2). This indicated the presence of two populations of growing hyphae in the BcCHSIIIa::eGFP strain, differing in apical fluorescence variability though time. In all the hyphae grouped in Cluster 2, the temporal fluorescence behavior at the apex was distinct from the one observed in hyphae of the control strain, suggesting a specific dynamic of SV in these hyphae.

Interestingly, a link between clustering and roundness of the apical fluorescent signal was noticed in hyphae in the BcCHSIIIa::eGFP strain. Cluster 1 grouped the hyphae with roundness values below 0.5 while Cluster 2 grouped the hyphae with roundness values above 0.5 (Fig.3B, Mann–Whitney U test, *p*-value 0.001). In comparison, neither hyphal width (Fig.3C) nor apex curvature (Fig.3D) significantly influenced the clustering outcome (Mann–Whitney U test, *p*-value 0.478, Student’s t-test, *p*-value 0.219, respectively). This indicates that the temporal dynamics which separate the acquisitions of Cluster 1 from those of Cluster 2 are associated with differences in the spatial organization of the SV-related apical fluorescence. On one hand, the hyphae belonging to Cluster 1 display a crescent accumulation of fluorescence at the apex (roundness < 0.5) whose variations over time could not be distinguished from the intrinsic fluorescence variations displayed by cytosolic eGFP. On the other hand, the hyphae belonging to Cluster 2 display a spherical accumulation of fluorescence at the apex (roundness > 0.5) whose variations over time statistically differed from the intrinsic dynamic of eGFP. This suggests that the measured variations of fluorescence in these hyphae originates from significant fluctuations in vesicle accumulation at the apex over time. The spherical accumulation of fluorescence at the apex being consistent with the presence of a spitzenkörper^18–20,40^, we considered that the Cluster 2 hyphae displayed a spitzenkörper.

**Figure 3.**
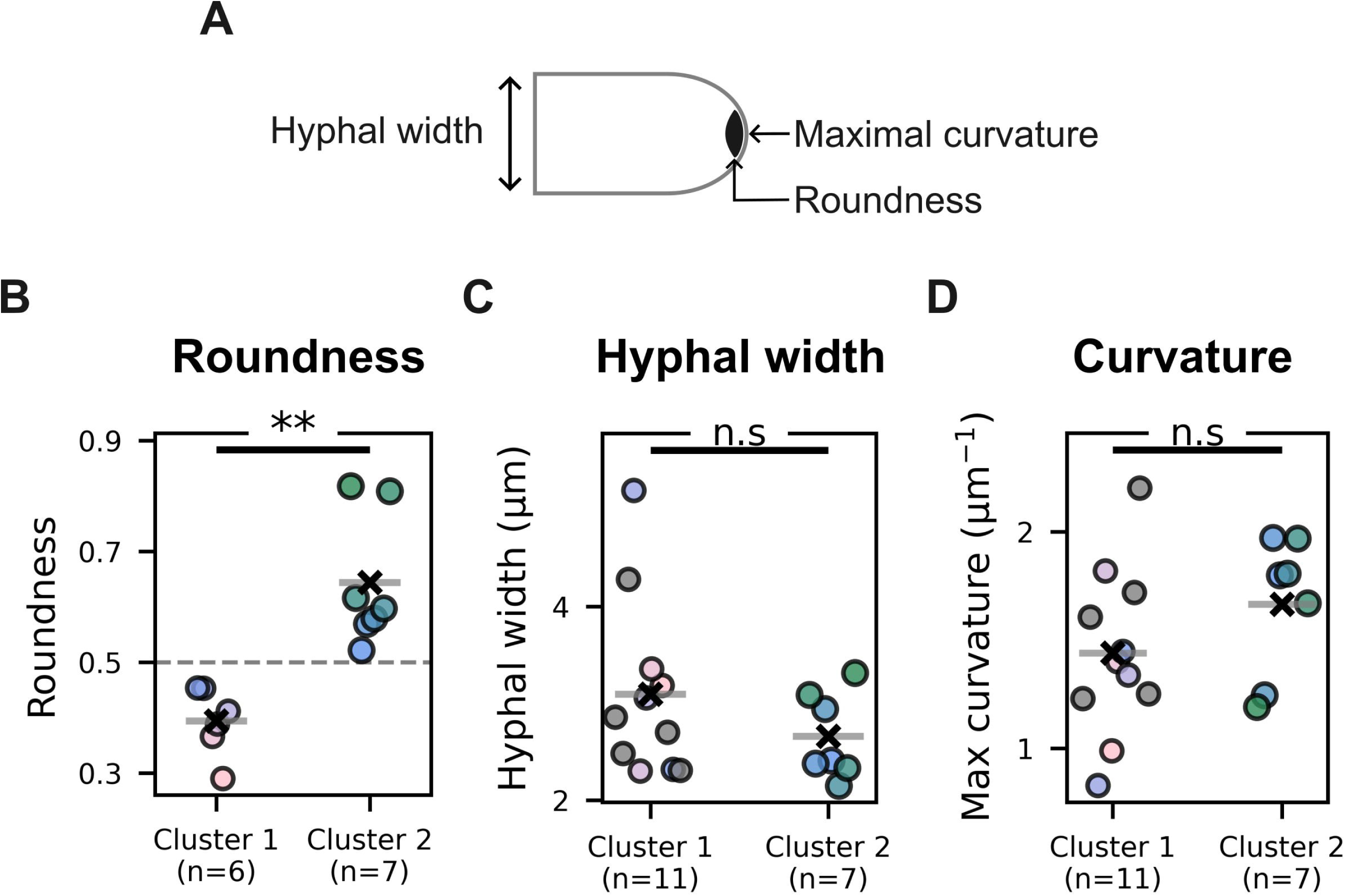
The roundness of the region of fluorescence accumulation differs between acquisitions belonging to the TDC Cluster 1 or Cluster 2. (A) Schematic representation of a hypha showing hyphal width, the point of maximal curvature and the region of fluorescence accumulation whose roundness was measured. (B) Roundness of the region of fluorescence accumulation measured on time projection of BcCHSIIIa::eGFP acquisitions, grouped by TDC clusters (Mann–Whitney U test, *p*-value 0.001). (C) Hyphal width measured for acquisitions of both control eGFP^Cyt^ and BcCHSIIIa::eGFP strains, grouped by cluster (Mann–Whitney U test *p*-value 0.479). (D) Maximum curvature of the apex grouped by cluster (Student’s t-test *p*-value 0.219). (B–D) Data points are color-coded according to the roundness measure, consistent with Figures 1 and 2. Data points of the eGFP^Cyt^ strain acquisitions are shown in gray.

### 5. Fluorescence variations are periodic in hyphae exhibiting a spitzenkörper

The production of kymographs from the time-lapse acquisitions of the BcCHSIIIa::eGFP strain revealed occurrences of fluorescence hotspots (Fig.S2), as exemplified by one acquisition (Hypha 12) shown in Figure 4A. These spots reflected spatially and temporally restricted events of fluorescence accumulation which appear as oscillations when the mean fluorescence intensity at the apex wasplotted against time (Fig.4B). Noticeably, these events appeared regularly spaced, suggesting the presence of an underlying periodic behavior.

**Figure 4.**
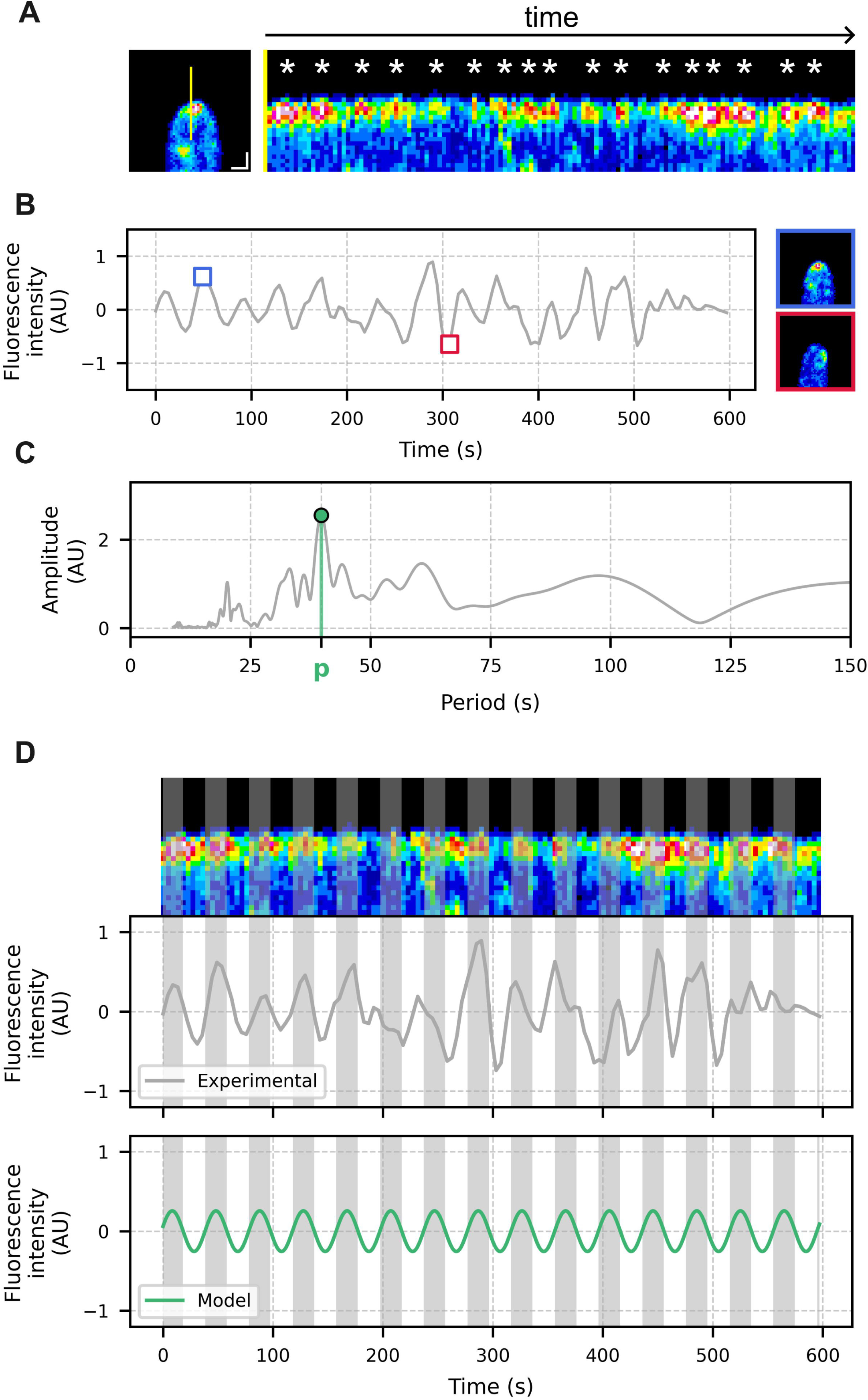
The spherical accumulation of fluorescence is marked by periodic variations of fluorescence. (A, left) Frame at t_0_ after tracking and cropping of the apex in the acquisition used as an example in this figure (Hypha 12). The yellow line indicates the position used to generate the kymograph. Scale bar = 1 µm. (A, right) Kymograph showing fluorescence intensity along the 3.8 µm line (yellow line in the left panel) over 135 frames (10 minutes total, 4.455 s interval). The absence of slope results from the dynamic tracking of the apex. White asterisks mark approximate positions of fluorescence hotspots. (B) Mean fluorescence intensity measured per frame over time. The blue square highlights a timepoint of high fluorescence and its corresponding frame (top right), the red square marks a low-intensity timepoint and its frame (bottom right). (C) Amplitude spectrum from Fast Fourier Transform analysis of the time series in (B), cropped to periods shorter than 150 seconds. The green marker indicates a peak corresponding to a period of 39.75 s.(D, top) Kymograph, (middle) the corresponding time series, (bottom) a sinusoidal model with a 39.75 s period. Grey zones represent predicted fluorescence peaks from the model and are overlaid on both the time series and the kymograph.

To evaluate the presence of a putative period governing the detected oscillations, the Fast Fourier Transform (FFT) was used. Pre-processed time series were analyzed (see *Material and methods*) and the outputs were expressed as amplitudes over periods, producing amplitude spectra (Fig.4C, Fig.5 left column). In the example shown in Figure 4C, the amplitude spectrum revealed a prominent oscillatory component with a period of 39.75 seconds. When this period was used to generate a sinusoidal model (Fig.4D, green line), this model closely aligned with the experimental data (Fig. 4D, grey line). Comparable results were obtained across all acquisitions grouped in Cluster 2 (Fig.5), each displaying a dominant oscillator with periods ranging from 31.39 to 64.93 seconds (mean period of 45.55 ± 10.09 seconds). All these acquisitions therefore shared a temporal dynamic marked by periodic events of fluorescence increases. In contrast, no such periodic behavior could be detected in the acquisitions grouped in Cluster 1. The existence of a linear correlation between the oscillation period and the hyphal elongation rate was explored but not found (Fig.S5, Pearson correlation, *p*-value 0.944). More acquisitions would however be needed to draw definitive conclusions regarding the existence of such a relationship.

**Figure 5.**
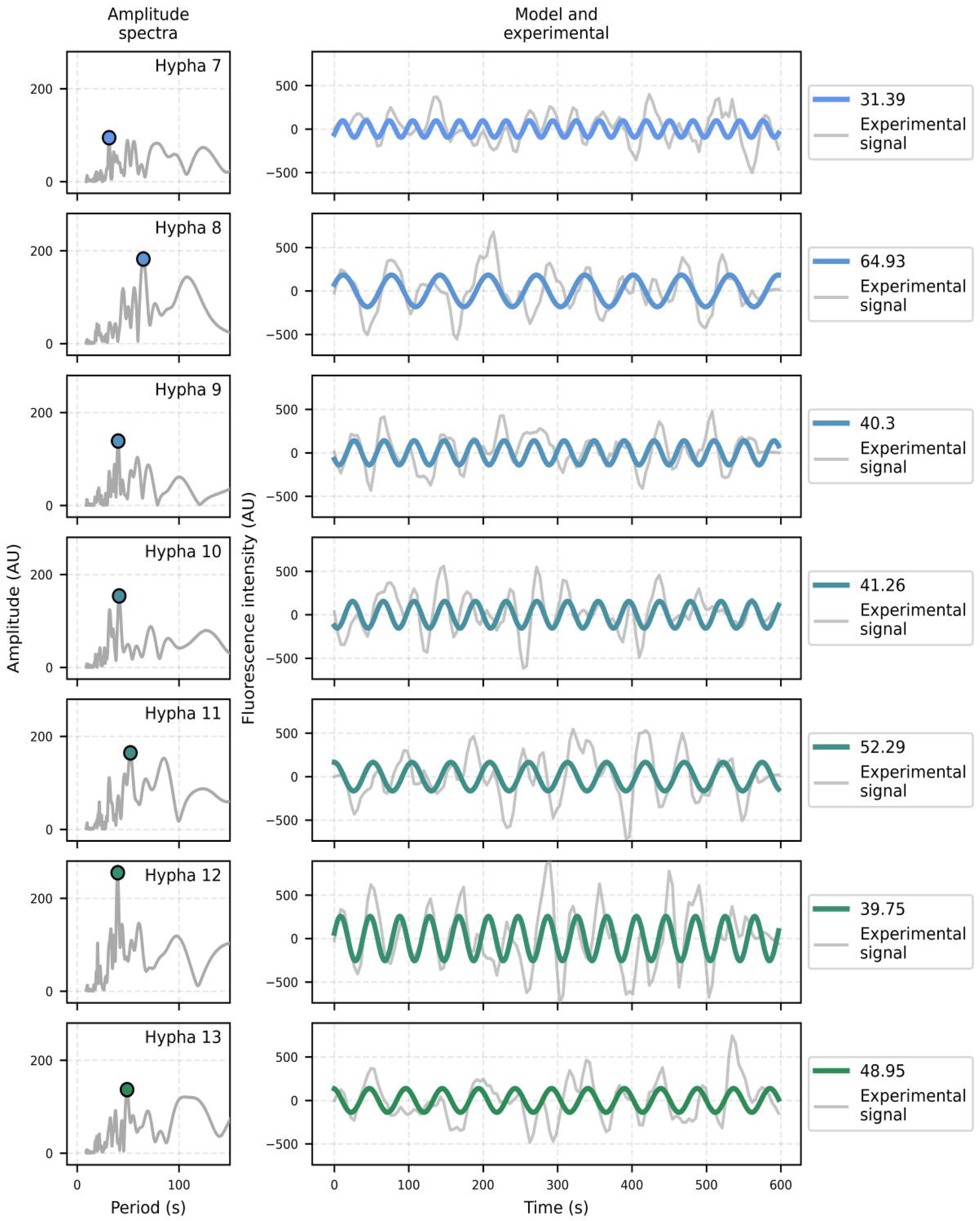
All hyphae assigned to Cluster 2 display a dominant oscillatory component. (Left column) Amplitude spectra obtained for acquisitions belonging to the TDC Cluster 2. Colored markers indicate the peak of maximal amplitude in each spectrum. For each acquisition, the peak amplitude and corresponding period were used to generate a sinusoidal model (colored bold line), overlaid on the experimental signal (grey line). Acquisitions are sorted in ascending order based on the roundness of the fluorescence accumulation region, such that the topmost acquisition has the lowest roundness value, and the bottommost has the highest. Models are color-coded according to the roundness measure, consistent with previous figures.

## Discussion

Polar cell elongation is a hallmark of vegetative growth in filamentous fungi. While the apical accumulation of Secretory Vesicles (SV) in a spitzenkörper has been broadly documented in Ascomycota and Basidiomycota, scarce observations of hyphae showing vesicle accumulation in a crescent^19,39,40^ suggested a diversity of SV accumulation patterns in growing hyphae. Here we used the fusion of a class III chitin synthase BcCHSIIIa (common SV marker) with the fluorescent protein eGFP to study the apical accumulation of SV in growing hyphae of the plant phytopathogen *B. cinerea*. As expected, we observed a concentrated accumulation of fluorescence at the apex of growing hyphae, but the shape of the region of fluorescence accumulation varied between hyphae, from spheroid to crescent-like. In line with the observation of a spitzenkörper in *B. cinerea* hyphae by electron microscopy^44^, the spheroid accumulation of fluorescence in some growing hyphae is consistent with the existence of a spitzenkörper in these hyphae, reflecting a spatially restricted accumulation of SV at their apex.

The crescent-shaped region of fluorescence accumulation reported here corresponds to an area of the apex within which the position of the recorded maximum fluorescence moves over time. Although less documented than accumulation at the spitzenkörper, a crescent organization was previously reported in *Aspergillus nidulans* expressing a fluorescently labeled class III chitin synthase^19^. It was also reported in the filamentous yeast *Ashbya gossypii* where SV and components of the exocyst and polarisome were labeled^39^. Altogether these observations suggest that a dual mode of apical SV accumulation can exist in filamentous fungi and may be shared among species. Our results document the existence of these two modes of SV distributions in growing hyphae of *B. cinerea*, but they also suggest that these two modes could be endpoints of a continuum rather than discrete categories, with intermediate forms distributed normally.

We observed a positive correlation between hyphal growth rate and the spatial organization of the fluorescence accumulating at the apex. Hyphae with fast elongation rates (>350 nm.min^-1^) consistently exhibited a spitzenkörper. This relationship is reminiscent of findings in *A. gossypii*, where the organization of the polarisome and exocyst, two complexes known to interact with the spitzenkörper, were shown to depend on growth rate, with faster-growing hyphae displaying a spherical organization and slower-growing ones showing a crescent-like arrangement^39^. Similarly, in *N. crassa*, immature hyphae lacking a spitzenkörper exhibit lower elongation rates than mature hyphae in which a spitzenkörper is present^33,45^. Altogether, these findings suggest a interdependence between spitzenkörper formation and hyphal elongation. However, whether vesicle accumulation facilitates faster growth or, conversely, rapid growth promotes spitzenkörper assembly remains to be elucidated. In addition, we also observed hyphae growing at similar rates (233 and 221 nm.min^-1^ for example) but exhibiting dissimilar apical SV organizations (crescent and spitzenkörper with roundness of 0.290 and 0.597, respectively in this example). Intermediate elongation rates may therefore be permissive to either the presence or absence of the spitzenkörper. Such a buffer zone was observed in *A. gossypii*, where some elongation rates were observed in hyphae with crescent or spitzenkörper-like organization of the exocyst or of the polarisome^39^. Altogether this suggests a not-strictly linear relationship between elongation rate and SV organization and these observations across species could support a model in which SV apical accumulation undergoes growth rate-dependent reorganization, transitioning from a crescent to a spitzenkörper as the elongation rate increases. With none of our recordings capturing hyphae undergoing substantial stable change in elongation rate, such a transition could not be observed. Whether such a reorganization occurs progressively within individual hyphae or whether crescent and spitzenkörper represent distinct and stable states associated with different hyphal populations therefore remains an open question.

We next investigated the temporal dynamics of apical fluorescence accumulation. Time-lapse confocal imaging revealed that fluorescence intensity fluctuated over time. To characterize these temporal patterns, we applied Temporal Dynamics Clustering^43^ (TDC) and this identified two distinct clusters: Cluster 1 grouping time series whose fluorescence profiles were indistinguishable from those of the control strain expressing cytosolic eGFP, and Cluster 2 grouping time series with marked fluctuations. Remarkably, all hyphae showing fluorescence accumulation in a crescent-shaped apical region grouped in Cluster 1, whereas hyphae with a spitzenkörper grouped in Cluster 2. Differences in the spatial organization of SV accumulation are therefore also associated with differences in temporal behavior (the fluorescence in hyphae displaying a spitzenkörper exhibit marked variations in intensity over time).

We further characterized the temporal dynamics present in acquisitions of hyphae displaying a spitzenkörper (Cluster 2) using the Fourier transform analysis. In all these hyphae, a dominant oscillatory component was found, indicating a repeated succession of high and low fluorescence intensity events regularly spaced over time. These variations are consistent with a periodic succession of SV accumulation and dispersion events at the apex and such periodic apical dynamics were previously described in *N. crassa* and *A. nidulans*^46^ as well as in *Arthrobotrys flagrans*^18^. Interestingly, the study by Takeshita *et al.*^46^ led to the proposition of an inverse relationship between the hyphal elongation rate and the oscillation period across species. In *N. crassa,* elongation is rapid (12 µm.min^-1^) and the period governing the apical fluorescent oscillation is short (17 ± 6 seconds)^46^. In *A. nidulans,* elongation is slower (1.8 µm.min^-1^) and the period is longer (30 ± 7 seconds)^46^. Our results in *B. cinerea* would support this trend with an even slower hyphal mean elongation rate (0.34 ± 0.21 µm.min^-1^) and a longer period governing the apical fluorescent oscillation (45 ± 10 seconds). While the methods used to identify this periodicity differ (measurement of the temporal distance between fluorescence peaks in the study of Takeshita *et al.*^46^ and Fourier transform analysis in our study), the observed pattern suggests that faster elongation rates may be associated with shorter oscillation periods at the species level. Such a relationship could be explained by the demand for lipids at the membrane level. Assuming stable size of SV, an increase in elongation rate would require an increase in lipid supply, made possible by a reduction in the time between exocytosis events. However, no such inverse relationship was observed when we compared periods and elongation rates between the 7 Cluster 2 hyphae of our study in *B. cinerea*. This absence may stem from insufficient sampling to detect subtler trends. A similar situation has been reported in pollen tubes, a tip-growing plant cell type that also exhibits periodic elongation, and in which a correlation between period and growth rate became apparent only with large sample sizes, due to high variability around the trend^47^.

Altogether, our study reveals the presence of two subpopulations in growing hyphae of *B. cinerea* that differ in the temporal dynamics of SV accumulation at the apex. One subpopulation shows relatively constant SV accumulation, while the other displays periodic fluctuations reflecting cycles of SV accumulation and release. This dichotomy was also observed in space, as the first subpopulation exhibited a crescent-shaped fluorescence signal, while the second showed a spherical accumulation reminiscent of the spitzenkörper. However, these two types of accumulation were not discrete classes but rather both ends of a continuum, revealing a plastic dimension to SV organization and accumulation in filamentous fungi. Recognizing this plasticity may reshape current models of fungal growth regulation, with broader implications for fungal development, adaptability, and pathogenicity. In light of these findings, it becomes interesting to investigate hyphal elongation in filamentous fungi with greater attention to the diversity of apical dynamics. Studies focusing solely on hyphae with a spitzenkörper may capture only a subset of fungal behavior and risk overlooking alternative mechanisms underlying polarized growth.

## Limitations of the study

While class III chitin synthases (CHS) are commonly used to visualize secretory vesicles via fusion with a fluorescent protein, it is important to note that these enzymes may be transported by only a subpopulation of vesicles. In *N. crassa*, CHS are specifically transported by microvesicles while the spitzenkörper also contains macrovesicles^31^. In *B. cinerea*, transmission electron microscopy has revealed the presence of both micro- and macrovesicles, but the distribution of BcCHSIIIa between these subpopulations remains unknown.

While the TDC approach did not identify fluorescence variations at the apex of the Cluster 1 hyphae (displaying a crescent-shaped region of fluorescence accumulation) when compared to the control strain hyphae, the existence of subtle variations cannot be ruled out. Acquisitions using more spatially and temporally resolutive microscopy are needed to further explore these dynamics.

The experimental design and confocal microscope limited the number of acquisitions. Additional data would help strengthen the analysis of the linear correlation between roundness of the region of accumulation and elongation rate. Nonetheless, statistical metrics and residual analysis provide a strong support for this relationship, even with the current sample size. Additional acquisitions of hyphae displaying a spitzenkörper will also help to uncover the relationship that connects period and elongation rate.

The signal pre-processing used in this study was intentionally stringent to minimize false positives, but it may have filtered out biologically informative signals, notably low-frequency patterns. Based on this work, we encourage future studies to further explore temporal dynamics that may have gone undetected in our study.

## Author contribution

AH conceived the study, performed the experiments, analyzed the data, developed the software tools, and prepared the figures. GC, MB, and M. Crumière produced the fungal strains used in this study. NP, M. Choquer, and CB secured funding and supervised the project. AH wrote the initial draft of the manuscript, which was reviewed and edited by AH, M. Choquer and CB, with input from all authors.

## Material and methods

### 1. Fungal strains

The strains BcCHSIIIa::eGFP and eGFP^Cyt^ were obtained by *Agrobacterium tumefasciens-* mediated transformation of the *B. cinerea* strain B05.10^48^. The BcCHSIIIa::eGFP strain was generated by the insertion, through homologous recombination, of an eGFP-tNiaD-NatR DNA fragment in frame with the BcCHSIIIa (Bcin04g03120) coding sequence (Fig.S1A). This DNA fragment, containing eGFP optimized for *B. cinerea*^42^, the terminator of the nitrate reductase from *B. cinerea* (tNiaD) and a nourseothricin resistance gene (NatR)^49^, was amplified from a donor plasmid. A 1031 bp fragment corresponding to the end of the BcCHSIIIa coding sequence, minus the STOP codon, was then amplified from B05.10 genomic DNA. A 924 bp fragment was amplified, corresponding to the 3’ UTR and terminator region of the BcCHSIIIa gene. The eGFP-tNiaD-NatR fragment was then assembled between the BcCHSIIIa coding sequence and the BcCHSIIIa terminator fragment using double-joint PCR^50^. This DNA cassette was inserted by *In Vivo* Assembly (IVA) cloning^51^ into a *A. tumefasciens* transfer-DNA carried by the p7 vector. Following verification by restriction enzyme digestion and sequencing, the vector was introduced in *A. tumefasciens* (strain LDA 1126) and the transformed bacteria was used to mutagenize *B. cinerea*^48^.

For the eGFP^Cyt^ strain, eGFP was placed under the control of the constitutive OliC promoter (pOliC) from *A. nidulans*^52^ and the tNiaD terminator. The pOliC-eGFP-tNiaD cassette was then inserted in the *A. tumefasciens* tDNA carried out by a pBHT2 plasmid containing a hygromycin resistance cassette^53^. Transformation led to the obtention of *A. tumefasciens* transformants that were later used to transform conidia of the parental B05.10 strain. *B. cinerea* transformants displaying hygromycin and nourseothricin resistance where then screened using epifluorescence microscopy. One transformant with strong signal was then selected for use in this study.

For all strains, conidia were harvested after 10 days of culture at 21°C with near-UV light on a medium containing 0.5% of glucose, 0.1% of malt extract, 0.1% of tryptone, 0.1% of casamino acids, 0.1% of yeast extract, 0.02% of ribonucleic acids and 1.6% of agar^54^. Conidia were then stored at -80°C and defrosted on ice before use in experiments.

### 2. Image acquisition

Before acquisition, conidia were diluted to 1.10^3^ conidia.ml^-1^ in MMII medium (NaNO_3_ 24 mM, Glucose 111 mM, KH_2_PO_4_ 0.147 mM, MgSO_4_ 0.039 mM, KCl 0.134 mM, FeSO_4_/7H_2_O 0.7 µM) to inoculate wells of a 96 wells SensoPlate microplate (GREINER) with 200µl per well. Incubation was performed at 21°C, in the dark. Time-lapse acquisitions (10 min) were conducted using a laser scanning confocal microscope (ZEISS Observer 7 equipped with an LSM 800) and a Plan-Apochromat 63x Oil DIC objective (ZEISS). Excitation was achieved using a 493 nm laser, and fluorescence emissions were detected within the range of 491 to 573 nm. The laser scanning was performed with an averaging of 16 on a single z-position. To ensure a uniform acquisition speed, the size of the acquired region was standardized (295x224 pixels) leading to an acquisition rate of 4.456 seconds per frame.

### 3. Tracking of apexes

To compensate for the elongation of filaments occurring during the acquisition, a custom FIJI^55^ macro was designed to move a region of interest (ROI) of 50x50 pixels (6.78x6.78 µm) through time. Each frame was then cropped to obtain a 50x50 time-lapse acquisition with the apex (4 first micrometers of the hyphae) centered at each frame.

### 4. Morphological measures

The curvature of the apex was measured on non-cropped acquisitions using the Kappa plugin^56^ in FIJI. A 10 µm curved was placed to follow the apical dome, the curvature was then estimated as the curvature of the most apical point on the first frame of each acquisition. The hyphal width was calculated as the average of three measures performed in the distal part of the apical region (3.3 ± 1.2 µm) on the first frame of each non-cropped acquisition.

To measure the geometry and area of maximal fluorescence intensity at the apex, a time-projection was first performed using the average intensity function in FIJI. Then, images were thresholded to only keep pixels in the 1% of highest intensity. The shape and area of the obtained structure was measured directly in FIJI.

Kymographs were produced using the KymographBuilder^57^ plugin (v.3.0.0) in FIJI. For growth speed estimation, kymographs were produced using all time points of the acquisition (135 total points with a time between frames of 4.456 seconds).

### 5. From acquisition to time series

Before measuring the intensity of fluorescence, two approaches were used to suppress background pixels that would bias the measure of fluorescence per frame: one to suppress extracellular pixels and one to suppress cytosolic background signals.

To limit the impact from extracellular pixels, a 50x50 pixels ROI was placed in the extracellular region uncropped acquisitions from both the BcCHSIIIa::eGFP and the eGFP^Cyt^ strains. The mean fluorescence intensity in this ROI was measured for all acquisition times and added to twice the associated standard deviation such as 95% of extracellular pixels have values bellow the set threshold^58^. All pixels with intensity values below this threshold were replaced by NaN values. The threshold was computed for each acquisition independently. Thresholded acquisitions were then cropped (see *Tracking of apexes*). Once cropped, the mean fluorescence intensity was measured at all time points of an acquisition to produce a time series (*i.e.* the mean fluorescence intensity over time). This approach restricts the measure of fluorescence intensity to the inside of the hyphae (termed *threshold technic 1* in this section).

To limit the impact from cytosolic background fluorescence present in the acquisitions of the BcCHSIIIa::eGFP strain, and thus limit the measure of pixel intensity to the regions of fluorescence accumulation, a 16x16 pixels ROI was placed in the sub-apical region (5 ± 2 µm bellow the hyphal tip) post-acquisition. The mean fluorescence intensity in this ROI was measured for the first 20 frames and added to twice the associated standard deviation such as 95% of background signals are smaller than the threshold value^58^. This threshold was used as an estimated of the intensity of the cytosolic background signal. All fluorescence values below this threshold were replaced by NaN values. Thresholded acquisitions were then cropped (see *Tracking of apexes*). Once cropped, the mean fluorescence intensity was measured at all time points of an acquisition to produce a time series. This approach restricts the measure of fluorescence intensity to regions of strong fluorescence (termed *threshold technic 2* in this section).

Following both approaches, the mean pixel intensity was measured at each frame producing one time series per acquisition corresponding to the mean fluorescence intensity over time. For each signal (*i.e.* time series) a trend was identified using a Savitzky-Golay filter of window length 10 (polynomial degree of 2). Each signal was then subtracted by its own trend to obtain detrended signals. No normalization was applied to preserve the relative amplitude of short-term fluctuations.

### 6. Temporal Dynamics Clustering

Temporal Dynamics Clustering (TDC) is a featured-based clustering approach for time series^43^. This approach was performed on detrended signals obtained after the threshold technic 1 (see *From acquisition to time series*). Following the approach detailed by Shuyang Li *et al.*^43^, the first order difference was computed independently for all signals such as:

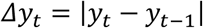

In our context, Δy_t_ therefore corresponds to the variation of fluorescence intensity between two successive frames. The second step of the TDC is the computation of *u* and of the subsequent *I_q_*, with:

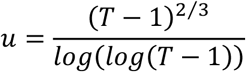

and:

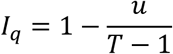

With T the length of the original time series. In our study, the number of time points is content between acquisitions leading to a single *u* and *I_q_* values for all time series. *I_q_* is a quantile index allowing the spreading of a time series into a *main* section containing all *y_t_* values below the and at the quantile *I_q_* and a *tail* section containing all *y_t_* values above *I_q_*. Following this partition of first-order differences, the *main* section contains moderate variations of fluorescence intensities while the *tail* section contains extreme events of fluorescence variation. These *main* and *tail* sections allow the production of metrics µ_main_ and µ_tail_ corresponding to the mean fluorescence intensities of the *main* and *tail* section respectively, and of σ_main_ that is the standard deviation of the *main* section. The values of µ_main_, µ_tail_ and σ_main_ are features used for the clustering using k-means clustering^59^. For a number of clusters from 2 to T-1, silhouette scores were computed to measure cluster quality and determine the optimal number of clusters, defined as the highest silhouette score with the minimal number of clusters.

### 7. Fast Fourier Transform

To study temporal variations in fluorescence intensity in regions of fluorescence accumulation, we further analyzed the detrended time series obtained after the threshold technic 2 (see section *From acquisition to time series*) using Fast Fourier Transform (FFT). Time series were first smoothed using a Savitzky-Golay filter (window size of 10, polynomial degree of 2) in combination with a moving average filter (window size of 3). To improve the frequency resolution, all signals were zero-padded to reach a total of 16 384 time points.

For each signal *S*, the FFT of *S* (FFT(S)) was computed for all discrete frequency bins *f* using the NumPy package^60^. The amplitude *A*(*f*) associated with each frequency *f* was computed as:

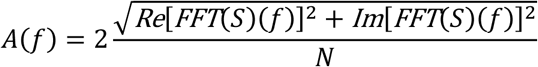

Where Re and Im denote the real and imaginary part of the FFT output, respectively, and N the number of points in the signal before zero padding. The factor 2 is used to correct the symmetrical distribution of the FFT output over positive and negative frequencies. For easier biological interpretation, amplitudes were plotted over periods (in seconds) instead of frequencies (in Hz). To satisfy the Nyquist sampling theorem, periods below 9 seconds were not considered. Additionally, only periods that could be sampled in at least four complete cycles during the 10-minute time-lapse acquisitions were considered, excluding periods longer than 150 seconds.

### 8. Sinusoidal model

For a period *p* and its associated amplitude *A_p_,* we created a sinusoidal model using the equation

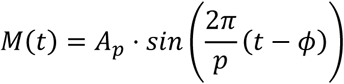

with ϕ the phase shift defined as

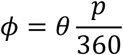

The value of ϕ was estimated as the value minimizing the Mean Squared Error between the model and the experimental data for values of θ between 0 and 360°.

### 9. Linear modeling and residual analysis

To produce linear models, the normality of the datasets was first evaluated using a Shapiro-Wilk test with a threshold *p*-value of 0.05. Peason and Spearman correlations were performed. The linear regression model was obtained by fitting an ordinal least squares regression using the statsmodels package^61^. A constant was added to the model to allow for a regression line with a non-zero intercept. Once the model was obtained, the 95% confidence interval was extracted using the statsmodels package and the R² computed. The influence of individual observations on the linear model was then assessed using residuals, standardized residuals and Cook’s distance. Observations with a Cook’s distance exceeding the 4/N threshold were considered influential outliers^62^.

### 10. Control for z-axis movement confirms genuine fluorescence dynamics

To rule out the possibility that the observed fluorescence variations resulted from z-axis movements during the recorded single-plane time-lapse acquisitions, we performed a time-lapse z-stack imaging on an hypha displaying a spitzenkörper (Fig.S4). The stack consisted of four focal planes (Z1 to Z4), with Z1 capturing the extracellular space and Z2–Z4 intersecting the fungal cell (Fig.S4A). First, time projection following apex tracking revealed a spheroid fluorescence accumulation in the 3 planes intersecting the fungal cell, with roundness values of 0.510, 0.832, and 0.575, respectively. Then, for each plane and frame, the mean intracellular fluorescence signals were extracted over time to produce time series (Fig.S4B). These signals were detrended to center them around zero (Fig.S4D) and finally compared through cross-correlation at lag 0. As shown in Figure S6E, the extracellular plane (Z1) and the intracellular planes (Z2-Z4) showed low similarity (correlation ≤ 0.3), whereas intracellular planes exhibited strong to very strong correlations (correlation = 0.6-0.9) between them. This indicates that the signals recorded from the 3 intracellular planes correspond to a single, consistent, intracellular fluorescence source. Artifacts caused by z-axis movements are therefore unlikely to cause the observed periodic fluorescence variations in the Cluster 2 hyphae.

### 11. Statistics

Throughout the article, measures are presented as mean ± standard deviation. When statistical tests were used, the name of the statistical test is indicated in parenthesis along with its associated *p* value. Statistic tests were performed in python using the scipy package^63^, with a significance threshold set at 0.05. In plots, ‘*ns’* indicates ‘non-significant’ tests (*p* > 0.05). A single asterisk (*) indicates *p* < 0.05, double asterisks (**) indicate *p* < 0.01 and triple asterisks (***) indicate *p* < 0.001.

## Supporting information

Figure S1. Construction of the BcCHSIIIa::eGFP strain

Figure S2. Kymographs obtained for all acquisitions of the BcCHSIIIa::eGFP strain

Figure S3. Validation metrics confirm linearity between roundness and growth rate

Figure S4. Cross-correlation analysis confirms consistent fluorescence oscillations across Z-positions

Figure S5. Periods of fluorescence variation are not linearly correlated with growth rate or the roundness of the region of fluorescence accumulation

Table 1. Quantitative measures for all analyzed acquisitions

## Figure legends

1. Figure S1. Construction of the BcCHSIIIa::eGFP strain (A) Schematic representation of the insertion of the *BcchsIIIa::eGFP-nat^R^-3’UTR* cassette by homologous recombination at the *BcchsIIIa* locus. A 1 031 bp fragment corresponding to the end of the *BcchsIIIa* coding sequence was fused with the coding sequence of the fluorescent protein eGFP, the *nat^R^* resistance cassette (conferring resistance to nourseothricin), and the 3’UTR of *BcchsIIIa*. (B) PCR amplification using primers AtMT_S-IIIa-GFP-RFP-F and AtMT_S_GFP-R confirms the presence of the genetic construct in the BcCHSIIIa::eGFP strain, and its absence in the parental strain. (C) Western-blot detection of the chimeric BcCHSIIIa::eGFP protein in the BcCHSIIIa::eGFP strain. The three lanes correspond to fractions obtained by differential centrifugation at 1 000g (first lane), 36 000g (second lane), and 100 000g (third lane), used to enrich secretory vesicles.
2. Figure S2. Kymographs obtained for all acquisitions of the BcCHSIIIa::eGFP strain Kymographs were generated along the elongation axis of individual hyphae to capture tip growth dynamics. For each acquisition, the fluorescence signal was measured over a 10 µm line at each time point, for a total duration of 10 minutes (135 time points with a 4.455-second interval). Each kymograph corresponds to one hypha. Kymographs are color-coded to represent fluorescence intensity. Scale bar = 5 µm. The table contains elongation rates as measures from kymographs for the 13 hyphae of the BcCHSIIIa::eGFP strain (Hypha 1 to Hypha 13) and the 5 hyphae of the eGFP^Cyt^ strain (Hypha 14 to 18). Measures are in nm.min^-1^.
3. Figure S3. Validation metrics confirm linearity between roundness and growth rate (A) Summary of key statistical metrics assessing the relationship between roundness and growth rate. Shapiro-Wilk tests support the normality of both distributions. (B, C) Q-Q plots for growth rate (B) and roundness (C). (D) Distribution of residuals from the linear model. (E) Q-Q plot of residuals. (F) Plot of standardized residuals versus fitted values. Orange and red lines indicate thresholds at ±2 and ±3, commonly used to identify moderate and extreme outliers, respectively. (G) Cook’s distance values used to identify influential points. Red line indicates the threshold 0.308 (calculated as 4/N with N=13).
4. Figure S4. Cross-correlation analysis confirms consistent fluorescence oscillations across Z-positions (A) Mean fluorescence intensity over time at the apex of a growing *Botrytis cinerea* hypha, obtained from the acquisition of time-lapse confocal images at four distinct Z-positions (Z1-Z4). (B) Schematic representation of the hypha and of focal planes. Z1 corresponds to a position outside the hypha, Z2 to Z4 are cross-sections of the cell. (C) Overview of the signal processing pipeline. (D) Processed fluorescence signals corresponding to the four Z-positions. (E) Cross-correlation matrix at lag 0 between processed signals. Black dashed box surrounds the region where the correlation score at lag0 is above 0.5.
5. Figure S5. Periods of fluorescence variation are not linearly correlated with growth rate or the roundness of the region of fluorescence accumulation (A) Scatter plot of period values measured in Cluster 2 acquisitions versus the elongation rate of the corresponding hyphae. (B) Scatter plot of period values measured in Cluster 2 acquisitions versus the roundness of the region of fluorescence accumulation. Points are numbered to match the corresponding hyphae shown in Figure 1C.

## Acknowledgement

We thank Isabelle Gonçalves for her critical reading of the manuscript. The authors also thank Bayer SAS for access to specific equipment.

## Declaration of interest

The authors declare that the research was conducted in the absence of any commercial or financial relationships that could be construed as a potential conflict of interest.

## Resource availability

For further information, contact Adrien Hamandjian. Data and code will be made available upon publication in a peer-reviewed journal.

---

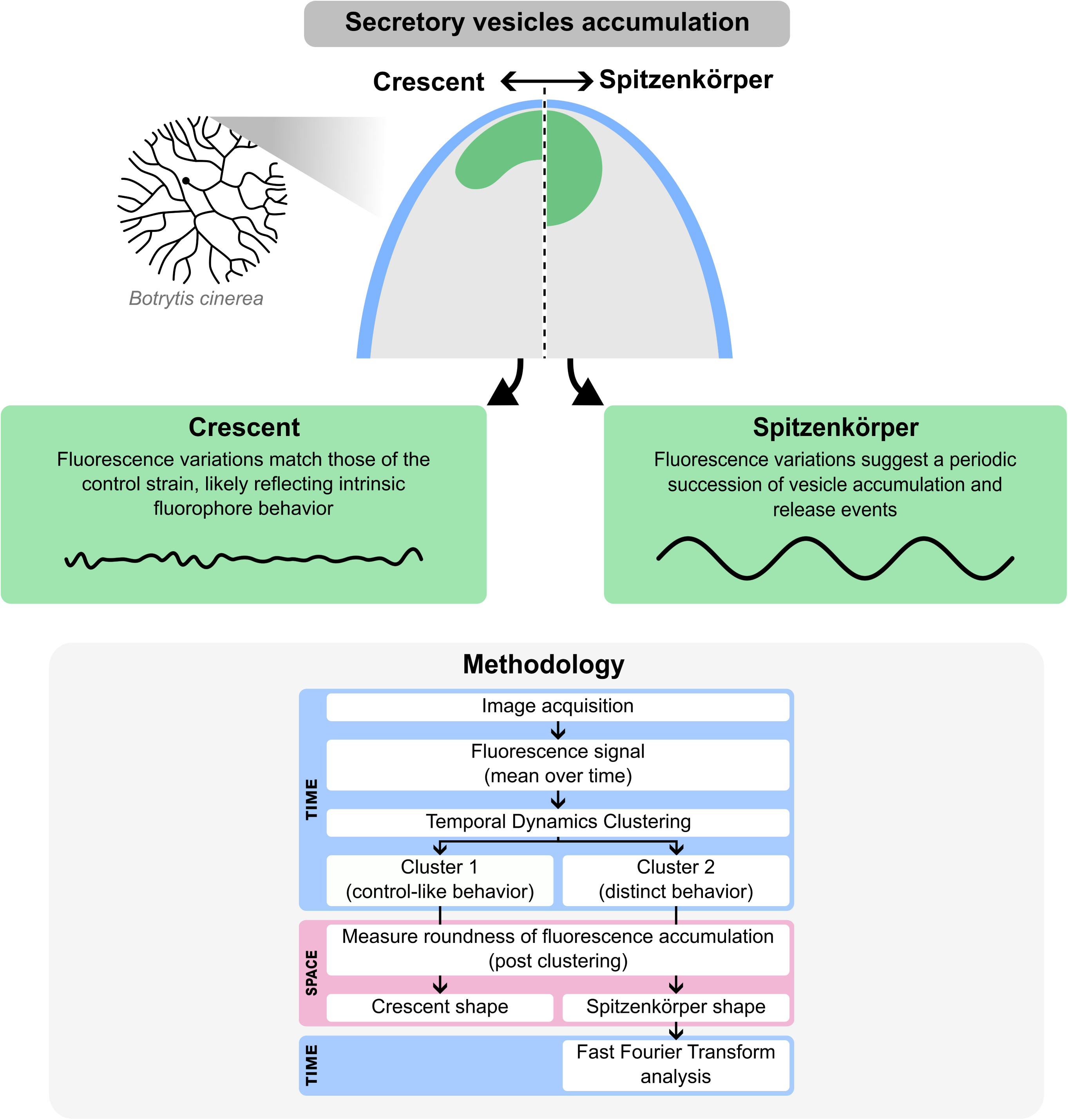

## Notes

### Competing Interest Statement

The authors have declared no competing interest.

### Summary of Updates

This revised version introduces major improvements in conceptual clarity, data interpretation, and manuscript structure. The Introduction was rewritten to more clearly frame the study around the diversity of apical secretory vesicle (SV) accumulation patterns in filamentous fungi. It now emphasizes how previous work has focused primarily on the spitzenkorper and sets up the need to re-examine polarized growth through the lens of SV geometry and dynamics. In the Results section, the distinction between the two modes of apical SV accumulation, spheroidal and crescent-shaped, was clarified and more precisely defined. Descriptions of signal localization and dynamics were rewritten to enhance accessibility. Quantitative comparisons between SV geometry and hyphal elongation rate were strengthened, and statistics were added to the linear model in addition to an analysis of residues. The normal distribution of the roundness is better introduced and explained. The temporal analysis of signal oscillations was clarified to be more accessible. The clustering approach and Fourier analysis are now better introduced, and the different steps are better explained. Descriptions of how different strains behaved were refined, and new interpretations of dynamic patterns were integrated into the main text. The section on the dynamics of elongation was taken out, while the approach was interesting the temporal and spatial resolution of our acquisitions was insufficient to provide interpretable results. Figures were updated accordingly to improve clarity, with cleaner labeling and the addition of summary plots and schema. The Discussion was substantially restructured. The revised version puts forward a more cohesive interpretation of the relationship between SV organization and growth dynamics. A major revision has been done to increase the clarity and ease the comprehension of the reader. The scientific Discussion was significantly increased. Additional hypotheses and concepts are introduced, notably the possibility of a dynamic continuum between crescent and spitzenkorper configurations. Existing hypotheses were reformulated to be clearer and more detailed. This version also more clearly states our results and acknowledges the limitations of the data. Additional comparisons with existing literature help contextualize the findings and refine the interpretation of polarity and vesicle organization in fungal cells. References were updated to include recent studies and provide a broader context. Across the manuscript, efforts were made to improve readability and reduce technical jargon, while preserving scientific depth. The overall structure and narrative were revised to better support the central argument and make the manuscript more accessible to a broad readership.

## References

1. Goriely, A., and Tabor, M. (2003). Biomechanical models of hyphal growth in actinomycetes. Journal of Theoretical Biology 222, 211–218. 10.1016/S0022-5193(03)00029-8.

2. Domozych, D.S., Fujimoto, C., and LaRue, T. (2013). Polar Expansion Dynamics in the Plant Kingdom: A Diverse and Multifunctional Journey on the Path to Pollen Tubes. Plants 2, 148–173. 10.3390/plants2010148.

3. Rensing, S.A. (2016). Plant Evo–Devo: How Tip Growth Evolved. Current Biology 26, R1228–R1230. 10.1016/j.cub.2016.09.060.

4. Boutillon, A., Banavar, S.P., and Campàs, O. (2024). Conserved physical mechanisms of cell and tissue elongation. Development 151, dev202687. 10.1242/dev.202687.

5. Braun, M., and Limbach, C. (2006). Rhizoids and protonemata of characean algae: model cells for research on polarized growth and plant gravity sensing. Protoplasma 229, 133–142. 10.1007/s00709-006-0208-9.

6. Mitchell, A.P. (1998). Dimorphism and virulence in *Candida albicans*. Current Opinion in Microbiology 1, 687–692. 10.1016/S1369-5274(98)80116-1.

7. Kadosh, D., and Johnson, A.D. (2005). Induction of the Candida albicans Filamentous Growth Program by Relief of Transcriptional Repression: A Genome-wide Analysis. Mol Biol Cell 16, 2903–2912. 10.1091/mbc.E05-01-0073.

8. Wendland, J. (2020). Sporulation in Ashbya gossypii. Journal of Fungi 6, 157. 10.3390/jof6030157.

9. Riquelme, M. (2013). Tip Growth in Filamentous Fungi: A Road Trip to the Apex. Annu. Rev. Microbiol. 67, 587–609. 10.1146/annurev-micro-092412-155652.

10. Peñalva, M.A., Zhang, J., Xiang, X., and Pantazopoulou, A. (2017). Transport of fungal RAB11 secretory vesicles involves myosin-5, dynein/dynactin/p25, and kinesin-1 and is independent of kinesin-3. Mol Biol Cell 28, 947–961. 10.1091/mbc.E16-08-0566.

11. Takeshita, N., Wernet, V., Tsuizaki, M., Grün, N., Hoshi, H., Ohta, A., Fischer, R., and Horiuchi, H. (2015). Transportation of Aspergillus nidulans Class III and V Chitin Synthases to the Hyphal Tips Depends on Conventional Kinesin. PLOS ONE 10, e0125937. 10.1371/journal.pone.0125937.

12. Mouriño-Pérez, R.R., Riquelme, M., Callejas-Negrete, O.A., and Galván-Mendoza, J.I. (2016). Microtubules and associated molecular motors in Neurospora crassa. Mycologia 108, 515–527. 10.3852/15-323.

13. Weiner, A., Orange, F., Lacas-Gervais, S., Rechav, K., Ghugtyal, V., Bassilana, M., and Arkowitz, R.A. (2019). On-site secretory vesicle delivery drives filamentous growth in the fungal pathogen Candida albicans. Cellular Microbiology 21, e12963. 10.1111/cmi.12963.

14. TerBush, D.R., Maurice, T., Roth, D., and Novick, P. (1996). The Exocyst is a multiprotein complex required for exocytosis in Saccharomyces cerevisiae. The EMBO journal, 15(23), 6483–6494.

15. Guo, W., Grant, A., and Novick, P. (1999). Exo84p Is an Exocyst Protein Essential for Secretion. Journal of Biological Chemistry 274, 23558–23564. 10.1074/jbc.274.33.23558.

16. Zuriegat, Q., Abubakar, Y.S., Wang, Z., Chen, M., and Zhang, J. (2024). Emerging Roles of Exocyst Complex in Fungi: A Review. Journal of Fungi 10, 614. 10.3390/jof10090614.

17. Gonçalves, I.R., Brouillet, S., Soulié, M.-C., Gribaldo, S., Sirven, C., Charron, N., Boccara, M., and Choquer, M. (2016). Genome-wide analyses of chitin synthases identify horizontal gene transfers towards bacteria and allow a robust and unifying classification into fungi. BMC Evol Biol 16, 252. 10.1186/s12862-016-0815-9.

18. Wernet, V., Kriegler, M., Kumpost, V., Mikut, R., Hilbert, L., and Fischer, R. (2023). Synchronization of oscillatory growth prepares fungal hyphae for fusion. eLife 12, e83310. 10.7554/eLife.83310.

19. Fukuda, K., Yamada, K., Deoka, K., Yamashita, S., Ohta, A., and Horiuchi, H. (2009). Class III Chitin Synthase ChsB of Aspergillus nidulans Localizes at the Sites of Polarized Cell Wall Synthesis and Is Required for Conidial Development. Eukaryot Cell 8, 945–956. 10.1128/EC.00326-08.

20. Sánchez-León, E., Verdín, J., Freitag, M., Roberson, R.W., Bartnicki-Garcia, S., and Riquelme, M. (2011). Traffic of Chitin Synthase 1 (CHS-1) to the Spitzenkörper and Developing Septa in Hyphae of Neurospora crassa: Actin Dependence and Evidence of Distinct Microvesicle Populations. Eukaryotic Cell 10, 683–695. 10.1128/EC.00280-10.

21. Viotti, C. (2016). ER to Golgi-Dependent Protein Secretion: The Conventional Pathway. In Unconventional Protein Secretion: Methods and Protocols, A. Pompa and F. De Marchis, eds. (Springer), pp. 3–29. 10.1007/978-1-4939-3804-9_1.

22. Girard, V., Dieryckx, C., Job, C., and Job, D. (2013). Secretomes: The fungal strike force. PROTEOMICS 13, 597–608. 10.1002/pmic.201200282.

23. Bani, A., Pioli, S., Ventura, M., Panzacchi, P., Borruso, L., Tognetti, R., Tonon, G., and Brusetti, L. (2018). The role of microbial community in the decomposition of leaf litter and deadwood. Applied Soil Ecology 126, 75–84. 10.1016/j.apsoil.2018.02.017.

24. de Vallée, A., Bally, P., Bruel, C., Chandat, L., Choquer, M., Dieryckx, C., Dupuy, J.W., Kaiser, S., Latorse, M.-P., Loisel, E., et al. (2019). A Similar Secretome Disturbance as a Hallmark of Non-pathogenic Botrytis cinerea ATMT-Mutants? Front Microbiol 10, 2829. 10.3389/fmicb.2019.02829.

25. Cairns, T.C., Zheng, X., Zheng, P., Sun, J., and Meyer, V. (2021). Turning Inside Out: Filamentous Fungal Secretion and Its Applications in Biotechnology, Agriculture, and the Clinic. J Fungi (Basel) 7, 535. 10.3390/jof7070535.

26. Riquelme, M., and Sánchez-León, E. (2014). The Spitzenkörper: a choreographer of fungal growth and morphogenesis. Curr Opin Microbiol 20, 27–33. 10.1016/j.mib.2014.04.003.

27. Gierz, G., and Bartnicki-garcia, S. (2001). A Three-Dimensional Model of Fungal Morphogenesis Based on the Vesicle Supply Center Concept. Journal of Theoretical Biology 208, 151–164. 10.1006/jtbi.2000.2209.

28. Egan, M.J., McClintock, M.A., and Reck-Peterson, S.L. (2012). Microtubule-based transport in filamentous fungi. Curr Opin Microbiol 15, 637–645. 10.1016/j.mib.2012.10.003.

29. Xie, Y., and Miao, Y. (2021). Polarisome assembly mediates actin remodeling during polarized yeast and fungal growth. Journal of Cell Science 134, jcs247916. 10.1242/jcs.247916.

30. Riquelme, M., Bredeweg, E.L., Callejas-Negrete, O., Roberson, R.W., Ludwig, S., Beltrán-Aguilar, A., Seiler, S., Novick, P., and Freitag, M. (2014). The Neurospora crassa exocyst complex tethers Spitzenkörper vesicles to the apical plasma membrane during polarized growth. Mol Biol Cell 25, 1312–1326. 10.1091/mbc.E13-06-0299.

31. Verdín, J., Bartnicki-Garcia, S., and Riquelme, M. (2009). Functional stratification of the Spitzenkörper of Neurospora crassa. Molecular Microbiology 74, 1044–1053. 10.1111/j.1365-2958.2009.06917.x.

32. Steinberg, G. (2007). Hyphal Growth: a Tale of Motors, Lipids, and the Spitzenkörper. Eukaryotic Cell 6, 351. 10.1128/EC.00381-06.

33. Araujo-Palomares, C.L., Castro-Longoria, E., and Riquelme, M. (2007). Ontogeny of the Spitzenkörper in germlings of Neurospora crassa. Fungal Genetics and Biology 44, 492–503. 10.1016/j.fgb.2006.10.004.

34. Schuster, M., Treitschke, S., Kilaru, S., Molloy, J., Harmer, N.J., and Steinberg, G. (2012). Myosin-5, kinesin-1 and myosin-17 cooperate in secretion of fungal chitin synthase. EMBO J 31, 214–227. 10.1038/emboj.2011.361.

35. Bracker, C.E., Murphy, D.J., and Lopez-Franco, R. (1997). Laser microbeam manipulation of cell morphogenesis growing in fungal hyphae. In Functional Imaging and Optical Manipulation of Living Cells (SPIE), pp. 67–80. 10.1117/12.274325.

36. Held, M., Kašpar, O., Edwards, C., and Nicolau, D.V. (2019). Intracellular mechanisms of fungal space searching in microenvironments. PNAS 116, 13543–13552. 10.1073/pnas.1816423116.

37. Thomson, D.D., Wehmeier, S., Byfield, F.J., Janmey, P.A., Caballero-Lima, D., Crossley, A., and Brand, A.C. (2015). Contact-induced apical asymmetry drives the thigmotropic responses of Candida albicans hyphae. Cell Microbiol 17, 342–354. 10.1111/cmi.12369.

38. Riquelme, M., Reynaga-Peña, C.G., Gierz, G., and Bartnicki-García, S. (1998). What Determines Growth Direction in Fungal Hyphae? Fungal Genetics and Biology 24, 101–109. 10.1006/fgbi.1998.1074.

39. Köhli, M., Galati, V., Boudier, K., Roberson, R.W., and Philippsen, P. (2008). Growth-speed-correlated localization of exocyst and polarisome components in growth zones of Ashbya gossypii hyphal tips. J Cell Sci 121, 3878–3889. 10.1242/jcs.033852.

40. Zhou, L., Evangelinos, M., Wernet, V., Eckert, A.F., Ishitsuka, Y., Fischer, R., Nienhaus, G.U., and Takeshita, N. (2018). Superresolution and pulse-chase imaging reveal the role of vesicle transport in polar growth of fungal cells. Sci Adv 4, e1701798. 10.1126/sciadv.1701798.

41. Soulié, M.-C., Perino, C., Piffeteau, A., Choquer, M., Malfatti, P., Cimerman, A., Kunz, C., Boccara, M., and Vidal-Cros, A. (2006). Botrytis cinerea virulence is drastically reduced after disruption of chitin synthase class III gene (Bcchs3a). Cellular Microbiology 8, 1310–1321. 10.1111/j.1462-5822.2006.00711.x.

42. Leroch, M., Mernke, D., Koppenhoefer, D., Schneider, P., Mosbach, A., Doehlemann, G., and Hahn, M. (2011). Living Colors in the Gray Mold Pathogen Botrytis cinerea: Codon-Optimized Genes Encoding Green Fluorescent Protein and mCherry, Which Exhibit Bright Fluorescence. Applied and Environmental Microbiology 77, 2887–2897. 10.1128/AEM.02644-10.

43. Li, S., Francini, G., and Magli, E. (2023). Temporal dynamics clustering for analyzing cell behavior in mobile networks. Computer Networks 223, 109578. 10.1016/j.comnet.2023.109578.

44. Roberson, R.W., Abril, M., Blackwell, M., Letcher, P., McLaughlin, D.J., Mouriño-Pérez, R., Riquelme, M., and Uchida, M. (2010). Hyphal structure, in cellular and molecular biology of filamentous fungi. Cellular and Molecular Biology of Filamentous Fungi, 8–24.

45. Guo, M., Kilaru, S., Schuster, M., Latz, M., and Steinberg, G. (2015). Fluorescent markers for the Spitzenkörper and exocytosis in *Zymoseptoria tritici*. Fungal Genetics and Biology 79, 158–165. 10.1016/j.fgb.2015.04.014.

46. Takeshita, N., Evangelinos, M., Zhou, L., Serizawa, T., Somera-Fajardo, R.A., Lu, L., Takaya, N., Nienhaus, G.U., and Fischer, R. (2017). Pulses of Ca2+ coordinate actin assembly and exocytosis for stepwise cell extension. Proceedings of the National Academy of Sciences 114, 5701–5706. 10.1073/pnas.1700204114.

47. Hemelryck, M.V., Bernal, R., Ispolatov, Y., and Dumais, J. (2018). Lily Pollen Tubes Pulse According to a Simple Spatial Oscillator. Sci Rep 8, 12135. 10.1038/s41598-018-30635-y.

48. Rolland, S., Jobic, C., Fèvre, M., and Bruel, C. (2003). Agrobacterium-mediated transformation of Botrytis cinerea, simple purification of monokaryotic transformants and rapid conidia-based identification of the transfer-DNA host genomic DNA flanking sequences. Curr Genet 44, 164–171. 10.1007/s00294-003-0438-8.

49. Crumière, M., de Vallée, A., Rascle, C., Gillet, F.-X., Nahar, S., van Kan, J.A.L., Bruel, C., Poussereau, N., and Choquer, M. (2025). A LysM Effector Mediates Adhesion and Plant Immunity Suppression in the Necrotrophic Fungus Botrytis cinerea. Journal of Basic Microbiology 65, e2400552. 10.1002/jobm.202400552.

50. Yu, J.-H., Hamari, Z., Han, K.-H., Seo, J.-A., Reyes-Domínguez, Y., and Scazzocchio, C. (2004). Double-joint PCR: a PCR-based molecular tool for gene manipulations in filamentous fungi. Fungal Genet Biol 41, 973–981. 10.1016/j.fgb.2004.08.001.

51. García-Nafría, J., Watson, J.F., and Greger, I.H. (2016). IVA cloning: A single-tube universal cloning system exploiting bacterial In Vivo Assembly. Scientific Reports 6, 27459. 10.1038/srep27459.

52. Souibgui, E., Bruel, C., Choquer, M., de Vallée, A., Dieryckx, C., Dupuy, J.W., Latorse, M.-P., Rascle, C., and Poussereau, N. (2021). Clathrin Is Important for Virulence Factors Delivery in the Necrotrophic Fungus Botrytis cinerea. Frontiers in Plant Science 12.

53. Mullins, E.D., Chen, X., Romaine, P., Raina, R., Geiser, D.M., and Kang, S. (2001). Agrobacterium-Mediated Transformation of Fusarium oxysporum: An Efficient Tool for Insertional Mutagenesis and Gene Transfer. Phytopathology 91, 173–180. 10.1094/PHYTO.2001.91.2.173.

54. Antal, Z., Rascle, C., Cimerman, A., Viaud, M., Billon-Grand, G., Choquer, M., and Bruel, C. (2012). The Homeobox BcHOX8 Gene in *Botrytis cinerea* Regulates Vegetative Growth and Morphology. PLOS ONE 7, e48134. 10.1371/journal.pone.0048134.

55. Schindelin, J., Arganda-Carreras, I., Frise, E., Kaynig, V., Longair, M., Pietzsch, T., Preibisch, S., Rueden, C., Saalfeld, S., Schmid, B., et al. (2012). Fiji: an open-source platform for biological-image analysis. Nat Methods 9, 676–682. 10.1038/nmeth.2019.

56. Mary, H., and Brouhard, G.J. (2019). Kappa (*κ*): Analysis of Curvature in Biological Image Data using B-splines. Preprint, 10.1101/852772.

57. Mary, H., Rueden, C., and Ferreira, T. (2016). KymographBuilder: Release 1.2.4. (Zenodo). 10.5281/zenodo.56702.

58. Ross, S.M. (2009). Chapter 2 - DESCRIPTIVE STATISTICS. In Introduction to Probability and Statistics for Engineers and Scientists (Fourth Edition), S. M. Ross, ed. (Academic Press), pp. 9–53. 10.1016/B978-0-12-370483-2.00007-2.

59. Pedregosa, F., Varoquaux, G., Gramfort, A., Michel, V., Thirion, B., Grisel, O., Blondel, M., Prettenhofer, P., Weiss, R., Dubourg, V., et al. (2011). Scikit-learn: Machine Learning in Python. Journal of Machine Learning Research 12, 2825–2830.

60. Harris, C.R., Millman, K.J., van der Walt, S.J., Gommers, R., Virtanen, P., Cournapeau, D., Wieser, E., Taylor, J., Berg, S., Smith, N.J., et al. (2020). Array programming with NumPy. Nature 585, 357–362. 10.1038/s41586-020-2649-2.

61. Seabold, S., and Perktold, J. (2010). Statsmodels: Econometric and Statistical Modeling with Python. scipy. 10.25080/Majora-92bf1922-011.

62. Altman, N., and Krzywinski, M. (2016). Analyzing outliers: influential or nuisance? Nature Methods 13, 281–282. 10.1038/nmeth.3812.

63. Virtanen, P., Gommers, R., Oliphant, T.E., Haberland, M., Reddy, T., Cournapeau, D., Burovski, E., Peterson, P., Weckesser, W., Bright, J., et al. (2020). SciPy 1.0: fundamental algorithms for scientific computing in Python. Nat Methods 17, 261–272. 10.1038/s41592-019-0686-2.

