## Supplementary figures and images for "Revisiting fungal polarized growth: from spitzenkörper to crescent accumulation of secretory vesicles"

### Figure S1. Construction of the BcCHSIIIa::eGFP strain

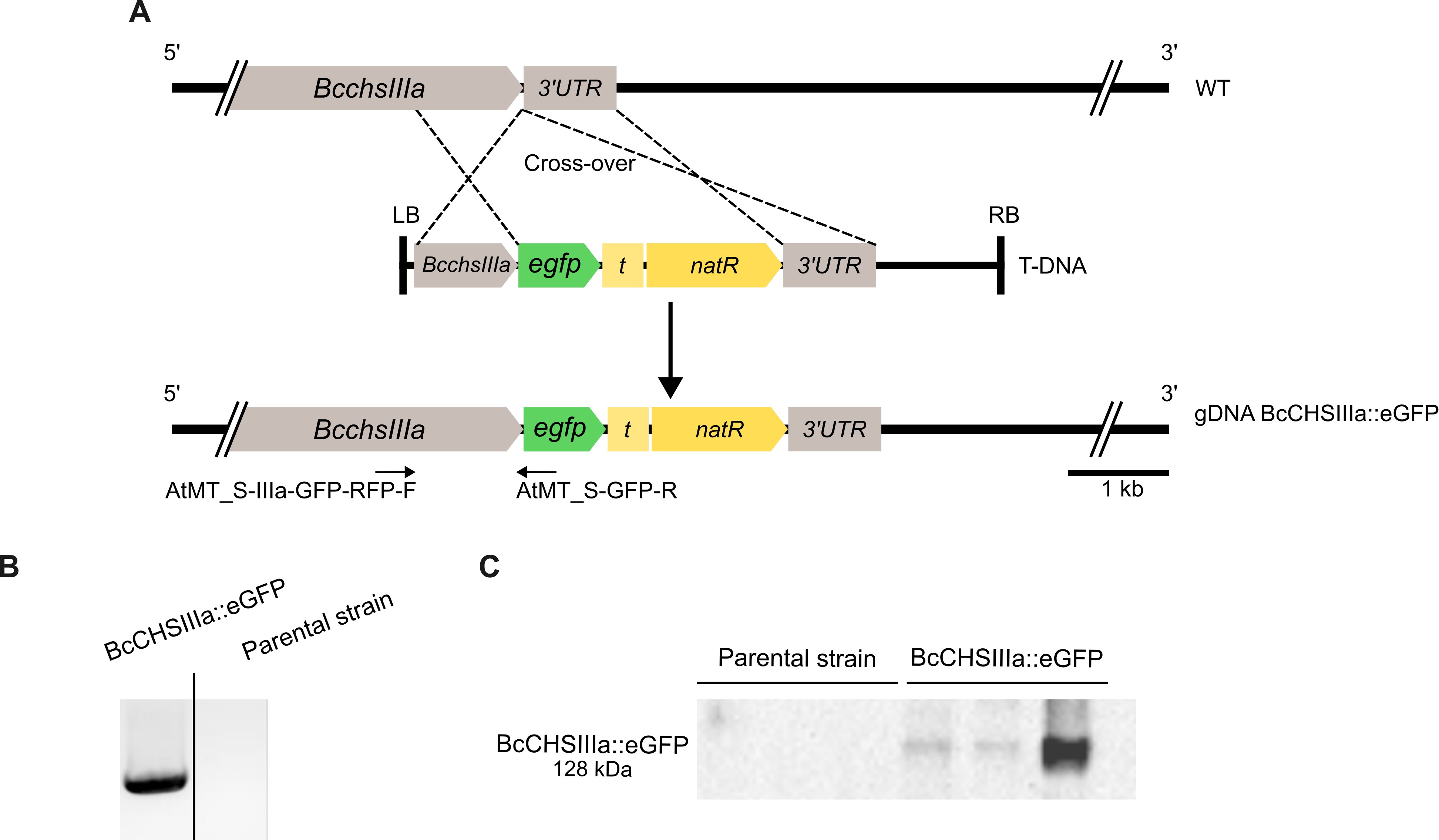

### Figure S2. Kymographs obtained for all acquisitions of the BcCHSIIIa::eGFP strain

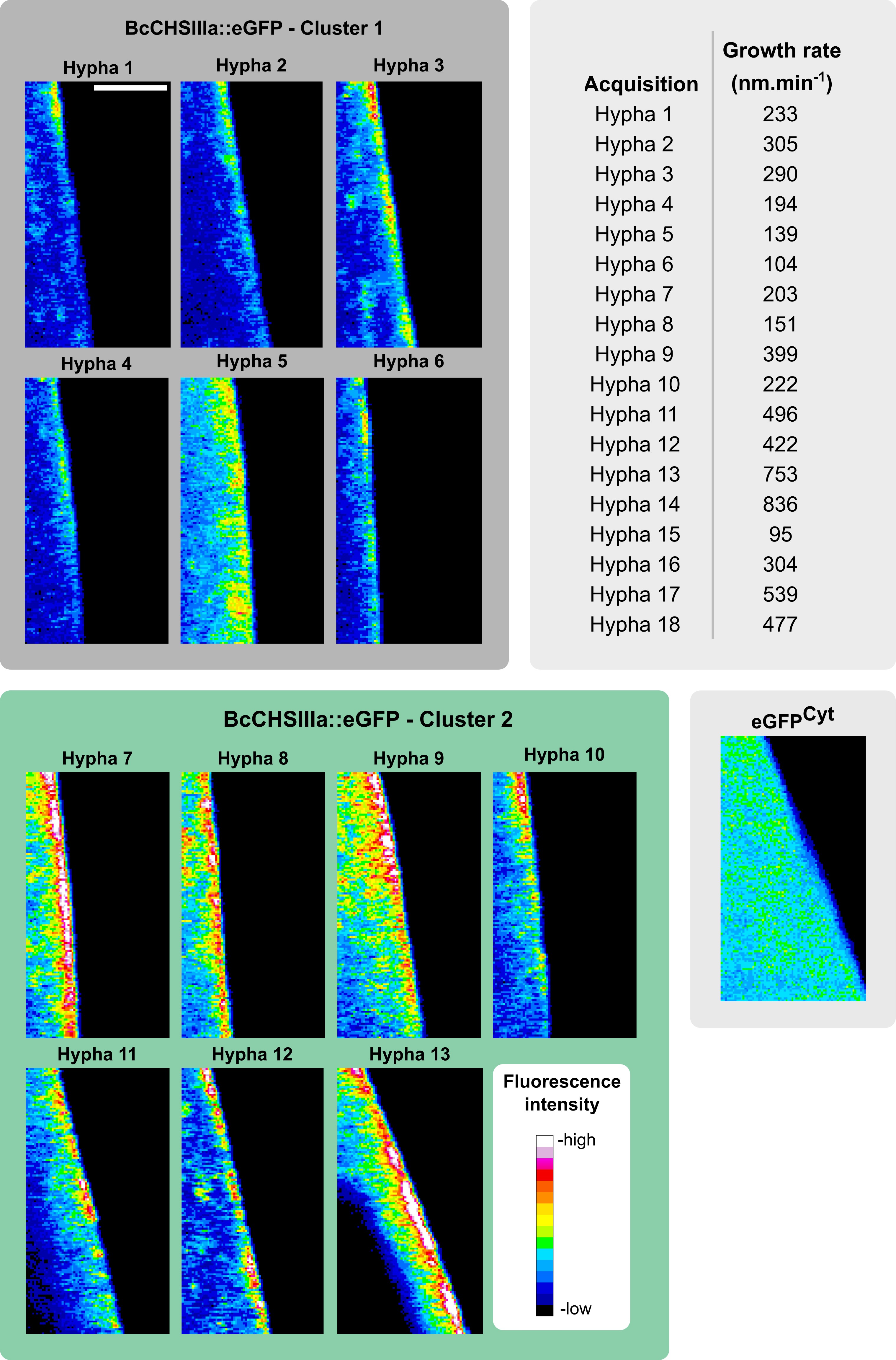

### Figure S3. Validation metrics confirm linearity between roundness and growth rate

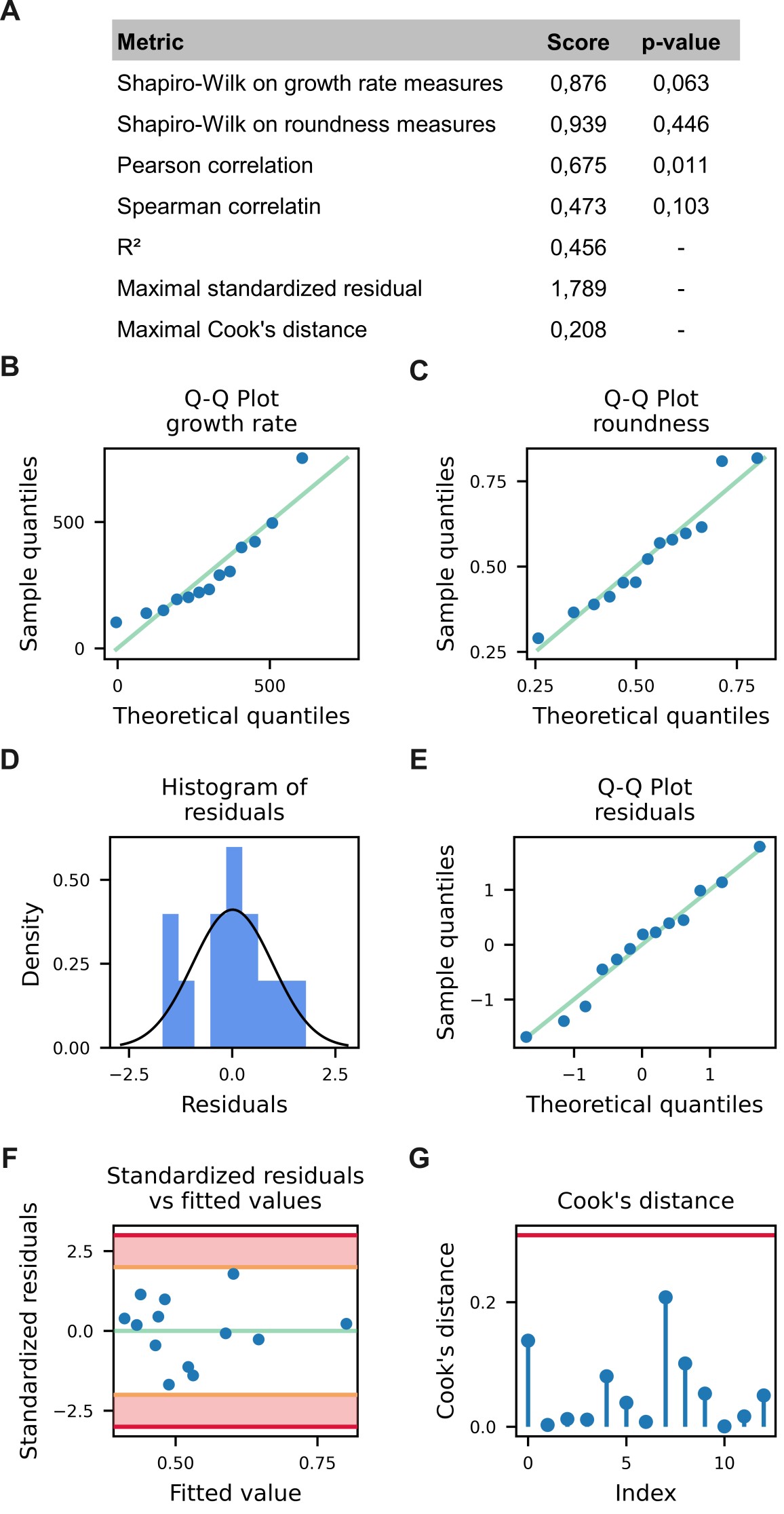

### Figure S4. Cross-correlation analysis confirms consistent fluorescence oscillations across Z-positions

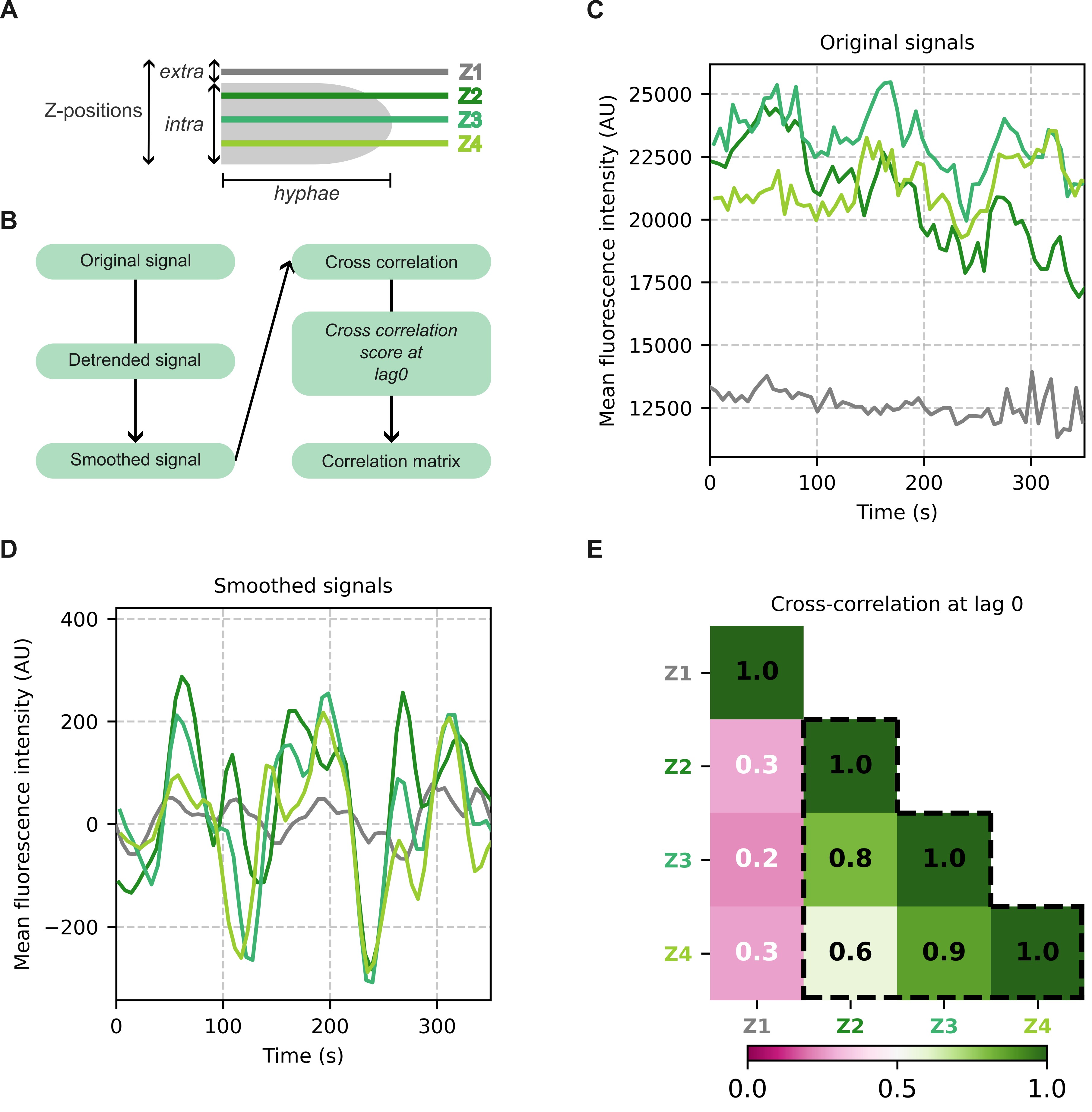

### Figure S5. Periods of fluorescence variation are not linearly correlated with growth rate or the roundness of the region of fluorescence accumulation

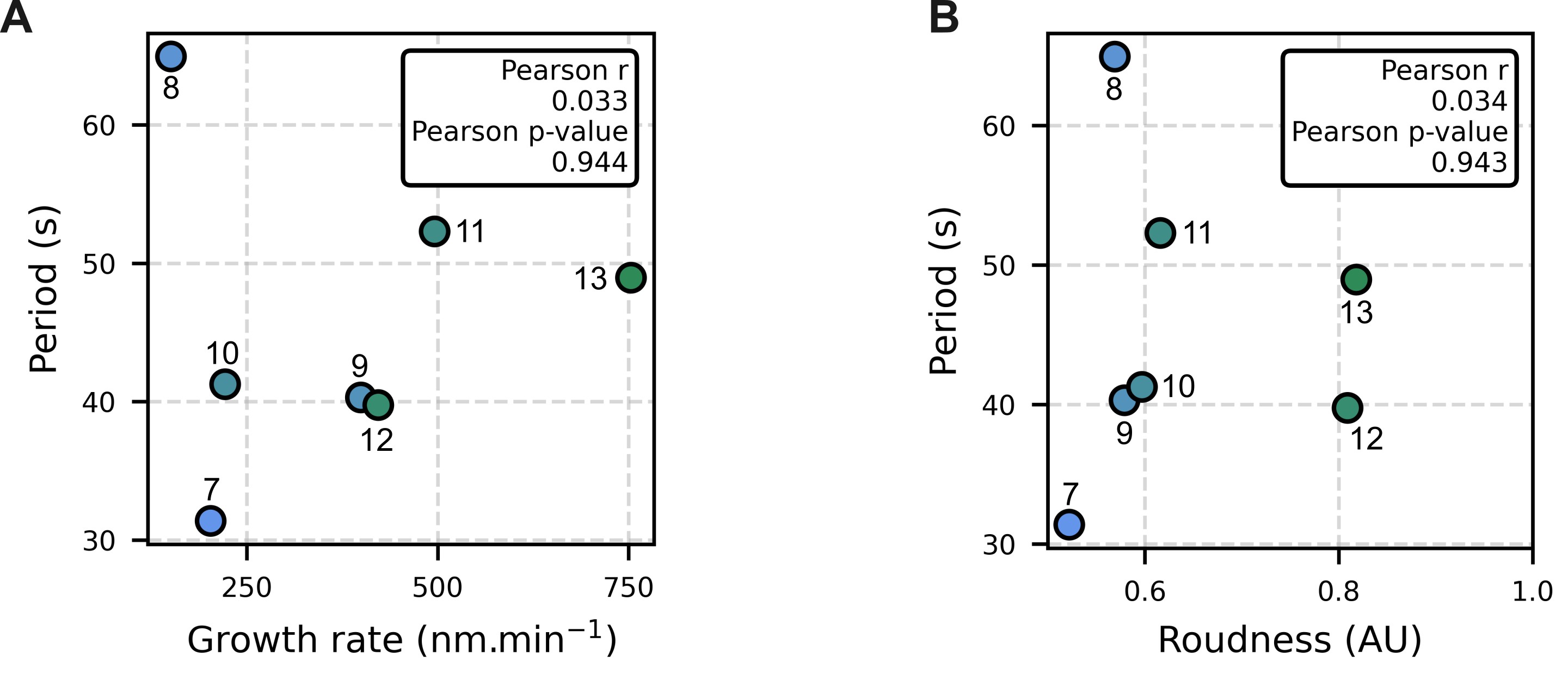

### Table 1. Quantitative measures for all analyzed acquisitions

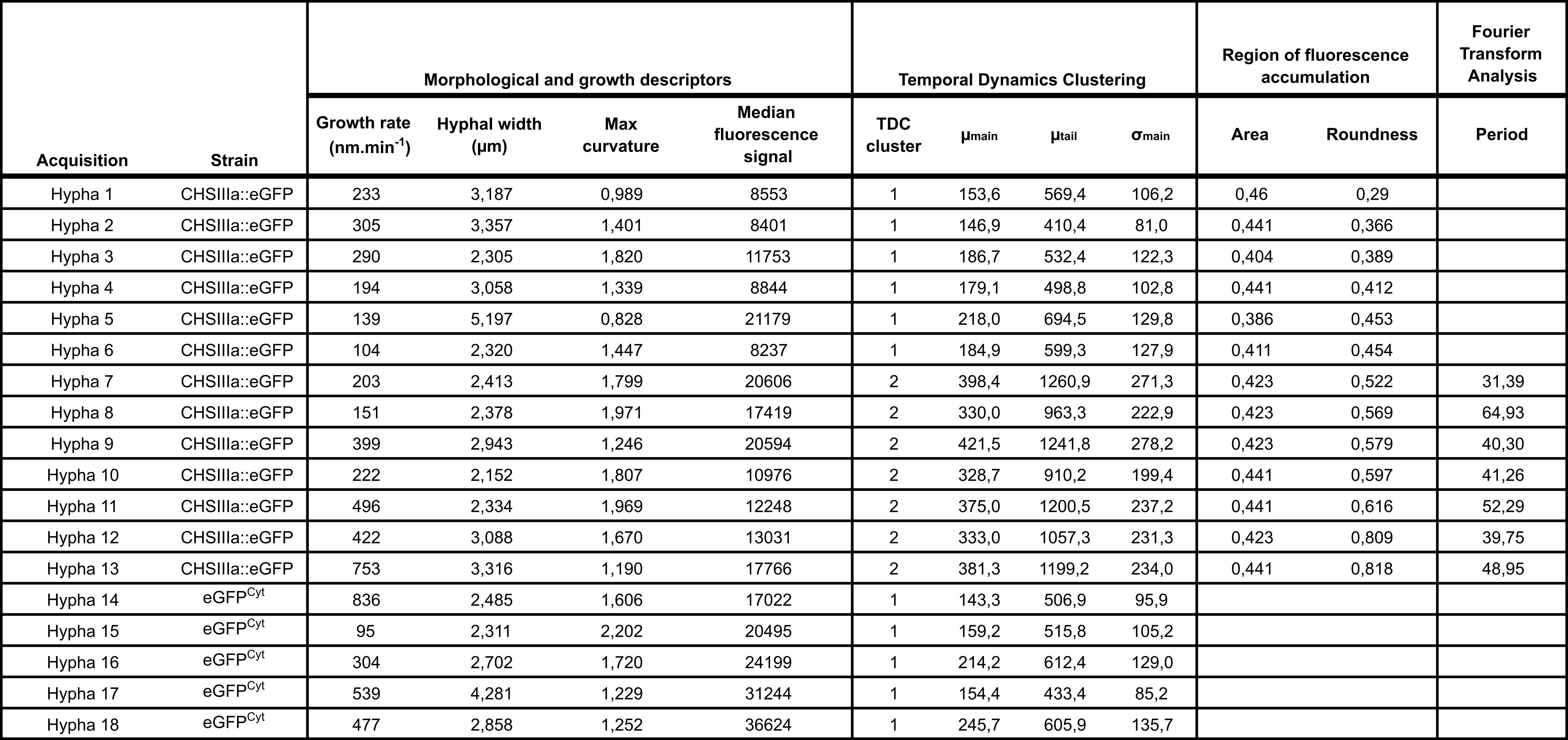
